# Highly Pathogenic Avian Influenza A (H5N1) clade 2.3.4.4b Virus detected in dairy cattle

**DOI:** 10.1101/2024.04.16.588916

**Authors:** Xiao Hu, Anugrah Saxena, Drew R. Magstadt, Phillip C. Gauger, Eric Burrough, Jianqiang Zhang, Chris Siepker, Marta Mainenti, Patrick J. Gorden, Paul Plummer, Ganwu Li

**Author notes:** Address for corresponding: Ganwu Li.

## Abstract

The global emergence of highly pathogenic avian influenza (HPAI) A (H5N1) clade 2.3.4.4b viruses poses a significant global public health threat. Until March 2024, no outbreaks of this virus clade had occurred in domestic cattle. We genetically characterize HPAI viruses from dairy cattle showing an abrupt drop in milk production. They share nearly identical genome sequences, forming a new genotype B3.13 within the 2.3.4.4b clade. B3.13 viruses underwent two reassortment events since 2023 and exhibit critical mutations in HA, M1, and NS genes but lack critical mutations in PB2 and PB1 genes, which enhance virulence or adaptation to mammals. The PB2 E627K mutation in a human case underscores the potential for rapid evolution post-infection, highlighting the need for continued surveillance to monitor public health threats.

## Introduction

Highly pathogenic avian influenza (HPAI) A (H5N1) virus belonging to clade 2.3.4.4b represents a significant global concern due to its severe impact on poultry populations, wildlife, and potential risks to human health. The origin of clade 2.3.4.4b can be traced back to 2020 when the H5N1 virus first emerged in domestic poultry in countries in East and Southeast Asia (https://wahis.woah.org#/home). Initial outbreaks were primarily confined to avian species, causing significant mortality among infected birds and posing substantial economic losses to the poultry industry (*1-3*). However, clade 2.3.4.4b exhibited a remarkable capability for geographic spread and host adaptation, leading to its dissemination across multiple continents through migratory bird pathways and global trade networks since 2020 (*3-8*). In late 2021, clade 2.3.4.4b H5N1 virus was introduced to North America from Eurasia and disseminated throughout the continent via wild birds, subsequently infecting numerous wild terrestrial mammals, such as foxes, skunks, bears, bobcats, and raccoons, posing a significant concern to public health (*5, 9, 10*).

Migratory birds play a significant role in transmitting HPAI viruses due to their ability to carry the virus over long distances (*11-14*). Texas lies within the Central Flyway, a major migratory flyway stretching from Canada to Mexico in North America (https://tpwd.texas.gov/huntwild/wild/birding/migration/flyways/). Additionally, Texas experiences some overlap in bird migration with neighboring states that belong to the Mississippi Flyway. This convergence of flyways heightens the risk of HPAI viral transmission, as migratory birds traverse diverse landscapes and habitats, including dairy cattle operations. In February and March 2024, a syndrome occurred in dairy cattle in the Texas panhandle region where affected animals developed a nonspecific illness and abrupt drop in milk production. Similar clinical cases were subsequently reported in dairy cattle in southwestern Kansas and northeastern New Mexico and mortalities in wild birds and domestic cats were observed within and around the affected sites in the Texas Panhandle. Here we present our findings on the detection, genomic characterization, phylogenetic analysis, and mutation adaptations of HPAI viruses, clade 2.3.4.4b H5N1, identified in dairy cattle, domestic cats, and wild birds in Texas. As this manuscript is being prepared for submission, the USDA has also confirmed the detection of this HPAI virus strain in dairy herds in Idaho, Michigan, Ohio, North Carolina, and South Dakota. Furthermore, the first human case of this virus in Texas, after contact with infected dairy cattle, has also been reported (https://www.dshs.texas.gov/news-alerts/dshs-reports-first-human-case-avian-influenza-texas).

## Results

### Detection of clade 2.3.4.4b HPAI viruses in domestic dairy cattle and cats in Texas in March, 2024

In February 2024, veterinarians in the Texas panhandle region observed lactating dairy cattle showing reduced feed intake, decreased milk production, and thickened yellow milk resembling colostrum. The syndrome peaked 4-6 days after onset and subsided within 10-14 days, mainly affecting older cows in mid to late lactation. By early March 2024, similar cases were reported in southwestern Kansas and northeastern New Mexico, with mortalities observed in wild birds and domestic cats near the affected areas. On March 21, 2024 milk samples from dairy cattle and fresh tissues from cats in Texas were received at the Iowa State University Veterinary Diagnostic Laboratory (ISU VDL). RT-PCR testing yielded positive results for influenza A virus (IAV) H5 clade 2.3.4.4b in the milk samples from the affected dairy cows along with brain and lung tissue from two domestic cats that reportedly consumed raw colostrum and milk at a dairy in Texas. The presence of highly pathogenic avian influenza (HPAI) H5N1 clade 2.3.4.4b was confirmed by National Veterinary Service Laboratories (NVSL) in Ames, IA, USA.

The two milk samples from two cows and brain and lung samples from two cats, which tested positive for IAV, underwent next-generation sequencing (NGS) and full genome sequences were successfully obtained on March 23, 2024 for subtyping and other further analyses. NGS analyses confirmed that all four individual samples were positive for HPAI A (H5N1). These sequences have been deposited in GenBank with the Bioproject number PRJNA1092030 (Supplementary Table 1). Several days later, the NVSL determined six HPAIV genome sequences from six wild birds, one sequence from a skunk, one from a human case, and an additional four from dairy cattle in Texas. These sequences are available in the GISAID database (https://gisaid.org) and were included in this study for analysis (Supplementary Table 2).

**Table 1.**
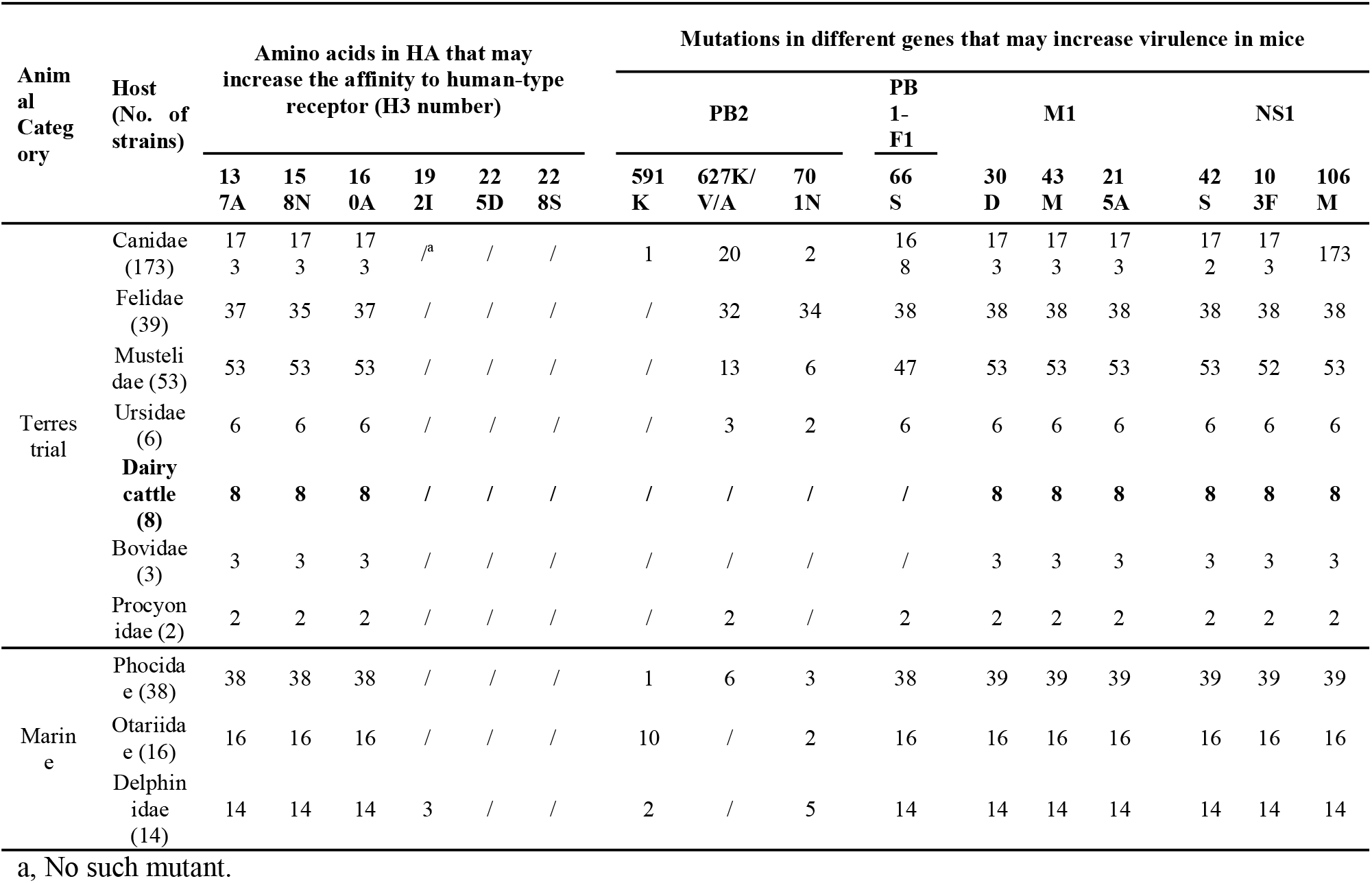
Mutations detected in the clade 2.3.4.4b H5N1 viruses have contributed to increased binding to human-type receptors and virulence in mammals.

### Phylogenetic and reassortment analysis

To track the genetically most closely related strains, we conducted a comprehensive search within the Global Initiative on Sharing Avian Influenza Data (GISAID) database. Supplementary Tables 3-10 present the results, indicating that the viruses isolated from wild birds, cows, cats, and humans in Texas during March 2024 shared a common ancestor with nearly 100 percent homology. To elucidate the phylogeny of HPAI A (H5N1) viruses in our study, we conducted individual phylogenetic analyses for each genome segment including a subset of HPAI A (H5N1) reference sequences obtained from avian and mammalian sources in America submitted to GISAID since January 1, 2021. Our analysis revealed that the genomes of viruses from two cows and two cats closely aligned and formed a cluster within the HPAI A (H5N1) subclade 2.3.4.4b shown in Figure 1 (for HA gene) and Supplementary Figures S1-S8 (for HA, NA, PB2, PB1, PA, NP, M, and NS genes). Specifically, we evaluated the time to the most recent common ancestor (tMRCA) of H5N1 viruses in Texas in 2024 by constructing the maximum clade credibility (MCC) tree of the HA gene using BEAST v1.10.4 (Figure 1). Referring to the lineage classification of clade 2.3.4.4b H5N1 viruses in the United States by GenoFlu (https://github.com/USDA-VS/GenoFlu) (*3*), the HA genes of the clade 2.3.4.4b H5N1 in America since 2021 were divided into three lineages: ea1, ea2, and ea3. The HA genes of our four HPAI H5N1 viruses, along with others from dairy cattle, wild birds, a skunk, and a human during this outbreak period in 2024, were grouped under lineage ea1. In addition, the NA genes were clustered within ea1, PB2 in am2.2, PB1 in am4, PA in ea1, NP in am8, M in ea1, and NS in am1.1 lineages (Figures S1-S8).

**Figure 1.**
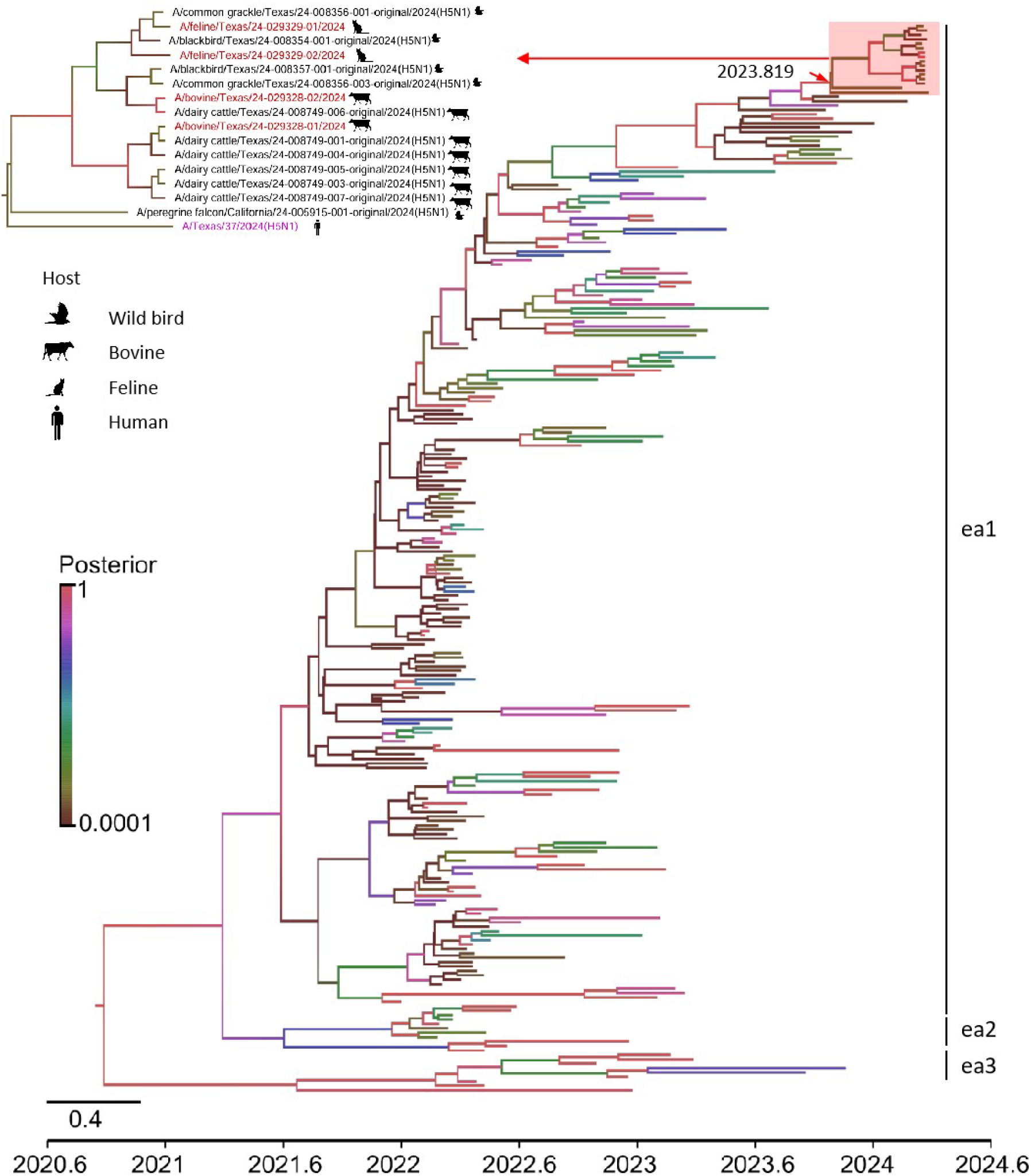
Maximum clade credibility (MCC) tree of the HA genes of clade 2.3.4.4b H5N1 viruses in the United States since 2021. The MCC tree is constructed by using BEAST v1.10.4 software package. Each branch is colored using posterior probability. The red frame represents H5N1 of Texas in 2024. The H5N1 viruses isolated in this study are shown in red, human isolate is shown in blue.

The automated data pipeline available at https://github.com/USDA-VS/GenoFlu was further applied to define their genotype (*3*). As illustrated in Figure 2, our four HPAI H5N1 viruses, along with others from dairy cattle, wild birds, a skunk, and a human during this outbreak period in 2024 belonged to genotype B3.13, resulting from a reassortment event involving genotype B3.7 and a low pathogenic avian influenza (LPAI) virus. The B3.7 genotype, which emerged in 2023, contributed seven gene segments, including PB2, PB1, PA, HA, NA, M, and NS, while the NP gene of B3.13 was originating from an LPAI virus resembling A/mallard/Alberta/567/2021 (11N9)-like strains. According to our GenoFlu analysis, the B3.7 genotype represents a 4+4 reassortant strain, with the HA, NA, PA, and MP genes originating from the H5N1 virus strain A1 in 2020, while the remaining segments (PB2, PB1, NP, and NS) are closely related to LPAI viruses. Our findings provide compelling evidence that the HPAI H5N1 viruses during this outbreak period in 2024 underwent reassortment events involving both HPAI and LPAI viruses.

**Figure 2.**
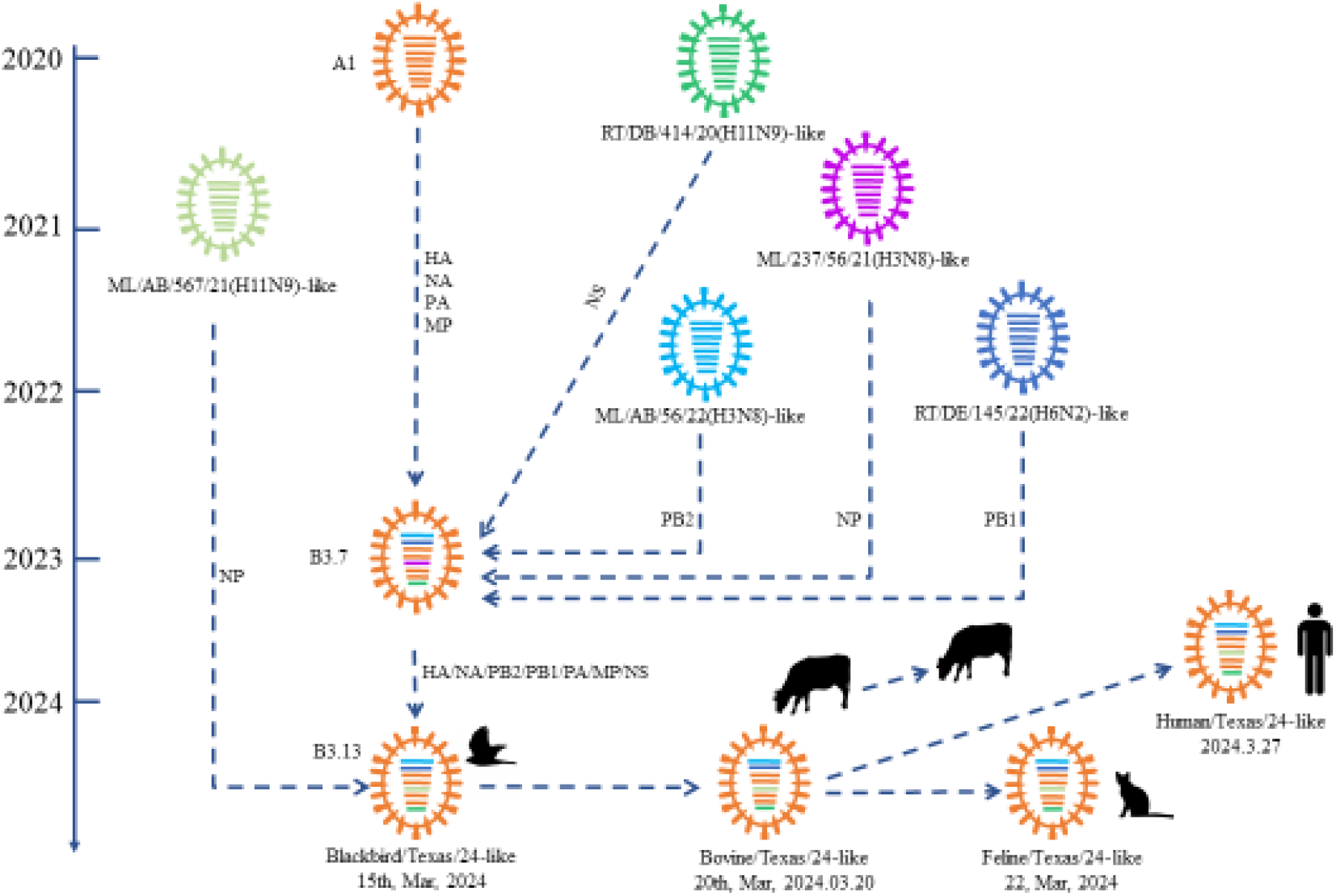
Schematic representation of genomic composition and reassortment time of HPAI H5N1 viruses from dairy cattle and other animals and human in March 2024. Viral particles are represented by colored ovals containing horizontal bars representing the eight gene segments (from top to bottom: PB2, PB1, PA, HA, NP, NA, M, and NS). Each color represents a separate virus background. The illustration is based on GenoFLU (https://github.com/USDA-VS/GenoFLU) and phylogenetic analysis. ML: mallard, RT: ruddy turnstone, AB: Alberta, DB: Delaware Bay.

### Critical amino acid mutation analysis

We conducted a comprehensive analysis of amino acid mutations, closely scrutinizing them to identify any changes potentially associated with increased affinity to human-type receptor heightened virulence, transmission or adaptation to mammalian hosts, and the mutants for antiviral resistance. We focused on comparing critical sites among eight HPAI virus isolates originating from dairy cattle and two from cats with those from terrestrial and marine mammals in the public source. This included an extensive dataset comprising 173 strains from Canidae, 39 strains from Felidae, 53 isolates from Mustelidae, six strains from Ursidae, three strains from other species of Bovidae, two strains from Procyonidae, as well as 68 marine mammal isolates, comprising 38 strains from Phocidae, 16 strains from Otariidae, and 14 strains from Delphinidae (Table 1). All 8 HPAI H5N1 isolates derived from dairy cattle and two cats demonstrated the presence of residues 137A, 158N, and 160A within their HA segments, which may increase binding affinity to the human-type receptor, while none contained residues 192I, 225D, or 228S (*15, 16*). This consistent pattern mirrors that observed in the majority of HPAI isolates from both terrestrial and marine mammals. Furthermore, all eight HPAI H5N1 isolates originating from dairy cattle and two cats exhibited residues 30D, 43M, and 215A in M1 (*17-19*), as well as 42S, 103F, and 106M in NS1 (*20*). Once again, this pattern is aligned with the prevalent composition observed across HPAI isolates from terrestrial and marine mammals, these mutants may increase the viral virulence in mammals. It is noteworthy that mutations 591K, 627K/V/A, or 701N in PB2, previously associated with mammalian host adaptation and enhanced transmission (*18, 21, 22*), were absent in all eight HPAI H5N1 isolates originating from dairy cattle and two cats, while the HPAI virus from the human case exhibited E627K mutation in PB2. Conversely, these mutations displayed a high frequency of occurrence in strains from Felidae and a lower frequency in strains from Canidae, Mustelidae, Phocidae, Otariidae, and Delphinidae.

Additionally, no critical site mutations associated with increased influenza antiviral resistance have been identified in the virus.

## Discussion

The widespread outbreaks of HPAI A (H5N1) clade 2.3.4.4b virus, since October 2020, have raised significant concerns regarding its impact on various mammalian species globally. Recent data reveal that, as of the latest assessment, 37 new mammal species have been afflicted since 2021. The majority of these cases involve wild terrestrial mammals such as foxes, skunks, bears, bobcats, and raccoons (*9, 23, 24*). Intriguingly, there have been sporadic infections among domestic pets like domestic cats and dogs (*25*), as well as marine mammals, including dolphins and sea lions (*26*). Moreover, from January 2022 to April 2023, eight documented human cases of H5N1 influenza from clade 2.3.4.4b have been recorded, several of which were severe or fatal (https://www.cdc.gov.flu/), underlining the gravity of this situation. Adding to this growing list of affected species, we now characterize an H5N1 influenza virus strain from clade 2.3.4.4b infecting dairy cattle associated with a sudden drop in milk production. The detection of this virus in bovine milk raises a potential public health concern related to zoonotic transmission through unpasteurized milk. This underscores the need for public awareness, pasteurization of milk to maintain adequate food safety, outbreak management, and a holistic approach to human health management.

In addition to being the first documented occurrence of HPAI A (H5N1) clade 2.3.4.4b virus infection in domestic dairy cattle, early pathology observations in this outbreak revealed an apparent tissue tropism for mammary gland in lactating domestic dairy cattle (personnel communication). Prior to this incident, the clade 2.3.4.4b IAV has typically caused systemic and respiratory diseases in wild mammals (*9*). Gross and microscopic lesions in wild mammals were frequently observed in organs such as the lung, heart, liver, spleen, and kidney, with some cases resulting in lesions in the brain leading to neurological signs. Furthermore, while it is widely recognized that certain strains of HPAI H5N1 clade 2.3.4.4b virus can breach the blood-brain barrier (*9, 23, 25, 27*), this is the first instance where the virus may penetrate the blood-milk barrier and be present in milk, raising potential public health concerns.

During this outbreak, HPAI virus strains from various sources such as wild birds, dairy cattle, cats, and a skunk, along with a human, displayed remarkably high nucleotide identities in their genome sequences, forming a distinct phylogenetic subcluster. These findings suggest an introduction of the 2.3.4.4b strain into Texas and neighboring regions by wild birds. The widespread detection of this HPAI virus strain across diverse regions and species underscores the complexity of its transmission pathways. Given the established role of migratory birds as reservoirs for avian influenza viruses (*11-14, 26, 28, 29*), it is important to highlight Texas’s location within the Central flyway. Texas also has overlap in bird migratory patterns with neighboring states that are part of the Mississippi Flyway. Furthermore, the outbreak’s occurrence in March coincides with the onset of the spring migration season, enhancing the likelihood of viral dissemination through migratory bird populations. Considering these factors, a highly plausible transmission route is hypothesized: wild birds may spread the virus through direct contact or contamination of water sources or feed staffs utilized by dairy cattle or other animals such as skunks. Consequently, other cattle in the herd, workers and domestic felids on dairy farms may contract the virus through direct contact with infected cattle or after consuming raw colostrum and milk from infected cattle. The detection of the same strain of HPAI viruses in various wild bird species, such as blackbirds and common grackles in Texas and Canada geese in Wyoming (Central Flyway), provides further support for this hypothesis. Another potential transmission scenario involves bovine-to-bovine spread. Recently, the USDA has verified the presence of this HPAI virus strain in dairy herds located in Idaho, Michigan, Ohio, North Carolina, and South Dakota (https://www.aphis.usda.gov/news/agency-announcements/usda-confirms-highly-pathogenic-avian-influenza-dairy-herd-idaho). In these cases, a documented history exists of cattle introduction from farms in the initial outbreak area, further supporting the hypothesis that lateral transmission can occur among cattle.

Our thorough examination of mutation adaptations, particularly those linked to human receptor binding affinity, increased virulence, transmission, or adaptation to mammalian hosts, offers critical insights into the risks posed by this specific strain of HPAI viruses. Notably, all HPAI viruses originating from dairy cattle and cats exhibit consistent amino acid residues in the HA gene, including 137A, 158N, and 160A, which have been documented to enhance the affinity of avian influenza viruses for human-type receptors (*15, 16*). Additionally, these dairy cattle-derived and cat-derived HPAI viruses harbor key virulence-increasing amino acid residues, such as 30D, 43M, and 215A in M1 (*17-19*), as well as 42S, 103F, and 106M in NS1(*20*). The presence of these amino acid mutations raises legitimate concerns regarding the potential for cross-species transmission to humans and other mammalian species. It is noteworthy that crucial mutations associated with mammalian host adaptation and enhanced transmission, specifically residues 591K, 627K/V/A, 701N, in PB2 (*18, 21, 22*), and 228S, along with the virulence-increasing residue 66S in PB1-F2(*30*), were conspicuously absent in all HPAI virus strains derived from dairy cattle and cats. This observation suggests that the current overall risk to human health is relatively low. However, it is imperative to recognize that influenza viruses have the capacity for rapid evolution within their host environments post-infection. A recent human case with direct contact with infected dairy cattle revealed a genetic change (PB2 E627K) (https://www.cdc.gov/flu/avianflu/spotlights/2023-2024/h5n1-analysis-texas.htm), indicating the potential for adaptation or transmission events. This underscores the dynamic nature of influenza viruses and the importance of continued surveillance and vigilance in monitoring potential threats to human health.

## Acknowledgement

We thank Dr. Amy Baker from USDA Agriculture Research Service for her critical review of this manuscript. Additionally, we extend our gratitude to the faculty and staff at the ISU VDL who contributed to the processing and analysis of clinical samples in this investigation.

## List of Supplementary Materials

Materials and Methods

Figures S1-S8

Table S1-S10

## Notes

### Competing Interest Statement

The authors have declared no competing interest.

## Reference

1. R. Xie et al., The episodic resurgence of highly pathogenic avian influenza H5 virus. Nature 622, 810–817 (2023).

2. N. S. Lewis et al., Emergence and spread of novel H5N8, H5N5 and H5N1 clade 2.3. 4.4 highly pathogenic avian influenza in 2020. Emerging Microbes & Infections 10, 148–151 (2021).

3. S. Youk et al., H5N1 highly pathogenic avian influenza clade 2.3. 4.4 b in wild and domestic birds: Introductions into the United States and reassortments, December 2021– April 2022. Virology 587, 109860 (2023).

4. F. A. Adlhoch C, Gonzales JL, Kuiken T, Marangon S, Niqueux É, et al, European Food Safety Authority; European Centre for Disease Prevention and Control; European Union Reference Laboratory for Avian Influenza. Avian influenza overview December 2021 - March 2022. 2022;20:e07289.

5. S. N. Bevins et al., Intercontinental movement of highly pathogenic avian influenza A (H5N1) clade 2.3. 4.4 virus to the United States, 2021. Emerging infectious diseases 28, 1006 (2022).

6. M. Agüero et al., Highly pathogenic avian influenza A (H5N1) virus infection in farmed minks, Spain, October 2022. Eurosurveillance 28, 2300001 (2023).

7. J. M. Rijks et al., Highly pathogenic avian influenza A (H5N1) virus in wild red foxes, the Netherlands, 2021. Emerging infectious diseases 27, 2960 (2021).

8. N. Ariyama et al., Highly pathogenic avian influenza a (H5N1) clade 2.3. 4.4 b virus in wild birds, Chile. Emerging Infectious Diseases 29, 1842 (2023).

9. E. J. Elsmo et al., Highly Pathogenic Avian Influenza A (H5N1) Virus Clade 2.3. 4.4 b Infections in Wild Terrestrial Mammals, United States, 2022. Emerging Infectious Diseases 29, 2451 (2023).

10. A. Kandeil et al., Rapid evolution of A (H5N1) influenza viruses after intercontinental spread to North America. Nature communications 14, 3082 (2023).

11. Y. Li et al., New avian influenza virus (H5N1) in wild birds, Qinghai, China. Emerging infectious diseases 17, 265 (2011).

12. M. Gilbert et al., Anatidae migration in the western Palearctic and spread of highly pathogenic avian influenza H5N1 virus. Emerging infectious diseases 12, 1650 (2006).

13. S. Zhou et al., Genetic evidence for avian influenza H5N1 viral transmission along the Black Sea–Mediterranean Flyway. Journal of General Virology 97, 2129–2134 (2016).

14. H. Tian et al., Avian influenza H5N1 viral and bird migration networks in Asia. Proceedings of the National Academy of Sciences 112, 172–177 (2015).

15. S. Yamada et al., Haemagglutinin mutations responsible for the binding of H5N1 influenza A viruses to human-type receptors. Nature 444, 378–382 (2006).

16. Y. Gao et al., Identification of amino acids in HA and PB2 critical for the transmission of H5N1 avian influenza viruses in a mammalian host. PLoS pathogens 5, e1000709 (2009).

17. S. Fan et al., Two amino acid residues in the matrix protein M1 contribute to the virulence difference of H5N1 avian influenza viruses in mice. Virology 384, 28–32 (2009).

18. A. Suttie et al., Inventory of molecular markers affecting biological characteristics of avian influenza A viruses. Virus Genes 55, 739–768 (2019).

19. N. Nao et al., A single amino acid in the M1 protein responsible for the different pathogenic potentials of H5N1 highly pathogenic avian influenza virus strains. PloS one 10, e0137989 (2015).

20. P. Jiao et al., A single-amino-acid substitution in the NS1 protein changes the pathogenicity of H5N1 avian influenza viruses in mice. Journal of virology 82, 1146–1154 (2008).

21. L. Bordes et al., Highly pathogenic avian influenza H5N1 virus infections in wild red foxes (Vulpes vulpes) show neurotropism and adaptive virus mutations. Microbiology spectrum 11, e02867–02822 (2023).

22. M. Hatta et al., Growth of H5N1 influenza A viruses in the upper respiratory tracts of mice. PLoS pathogens 3, e133 (2007).

23. B. D. Cronk et al., Infection and tissue distribution of highly pathogenic avian influenza A type H5N1 (clade 2.3. 4.4 b) in red fox kits (Vulpes vulpes). Emerging Microbes & Infections 12, 2249554 (2023).

24. T. Maemura et al., Characterization of highly pathogenic clade 2.3. 4.4 b H5N1 mink influenza viruses. EBioMedicine 97, (2023).

25. F. S. François-Xavier BriandComments to Author, Isabelle Pierre, Véronique Beven, Edouard Hirchaud, Fabrice Hérault, René Planel Angélina Rigaudeau, Sibylle Bernard-Stoecklin, Sylvie Van der Werf, Bruno Lina, Guillaume Gerbier, Nicolas Eterradossi, Audrey Schmitz, Eric Niqueux, and Béatrice Grasland, Highly Pathogenic Avian Influenza A(H5N1) Clade 2.3.4.4b Virus in Domestic Cat, France, 202. Emerging infectious diseases 29, (2023).

26. M. Leguia et al., Highly pathogenic avian influenza A (H5N1) in marine mammals and seabirds in Peru. Nature Communications 14, 5489 (2023).

27. L. Bauer, F. F. Benavides, E. J. V. Kroeze, E. de Wit, D. van Riel, The neuropathogenesis of highly pathogenic avian influenza H5Nx viruses in mammalian species including humans. Trends in Neurosciences, (2023).

28. G. Zhang et al., Bidirectional movement of emerging H5N8 avian influenza viruses between Europe and Asia via migratory birds since early 2020. Molecular Biology and Evolution 40, msad019 (2023).

29. M. Fourment, A. E. Darling, E. C. Holmes, The impact of migratory flyways on the spread of avian influenza virus in North America. BMC evolutionary biology 17, 1–12 (2017).

30. R. P. Kamal et al., Emergence of highly pathogenic avian influenza A (H5N1) virus PB1-F2 variants and their virulence in BALB/c mice. Journal of Virology 89, 5835–5846 (2015).

