## Supplementary materials for "Highly Pathogenic Avian Influenza A (H5N1) clade 2.3.4.4b Virus detected in dairy cattle"

**Domestic dairy cattle was detected by infecting with clade 2.3.4.4b highly pathogenic avian influenza A (H5N1) virus**

### Materials and Methods

#### Samples

Two bovine milk samples and fresh feline tissues from Texas were received at the ISU VDL on March 21, 2024 and were used in this investigation. These milk samples were from cattle reportedly exhibiting a nonspecific illness including reduced milk production.

#### Real-time RT-PCR and subtyping

Milk samples were diluted 1:4 v:v in phosphate buffered saline (PBS; pH 7.4) and tissues were processed into 10% homogenates in Earle’s balanced salt solution. Samples were extracted using the MagMAX^TM^ Viral RNA Isolation Kit and a Kingfisher Flex instrument. Reverse transcription real-time PCR (RT-rtPCR) was performed using the Vet MAX TM-Gold SIV Detection kit to screen for IAV RNA presence, with XENO internal positive control to monitor PCR inhibitors. Each RT-rtPCR plate included positive and negative controls. Positive samples (Ct values < 40.0) underwent H5 subtype and 2.3.4.4b clade H5 analysis using the same RNA extract and NAHLN approved RT-rtPCR. Reactions were conducted on an ABI 7500 Fast thermocycler with appropriate controls, considering samples with Ct values < 40.0 as positive.

#### Whole genome sequencing (WGS)

WGS was conducted on two milk samples and lung and brain tissue samples from two cats. Viral RNA was extracted using the MagMAX pathogen RNA kit and a KingFisher Flex system. Sequencing libraries were prepared using TruSeq. Next-generation sequencing was performed on an Illumina MiSeq platform following standard protocols at the ISU VDL. Approximately 2,000,000 raw sequencing reads per sample underwent preprocessing with Trimmomatic v0.36 and quality checking with FastQC. Quality-trimmed reads were aligned to reference sequences obtained from the NCBI Influenza Sequence Database using BWA-MEM. Mapped reads were then extracted using SAMtools and used for de novo assembly. Contigs for each segment were assembled using ABySS and SPAdes. Manual curation in SeqMan Pro was conducted to eliminate extraneous sequences and trim chimeric contigs, resulting in a consensus sequence for each segment.

#### Phylogenetic analysis

A maximum clade credibility (MCC) time-scaled phylogenetic tree of HA sequences from clade 2.3.4.4b H5N1 viruses was performed by Markov chain Monte Carlo (MCMC) method using BEAST v1.10.4. Reference sequences were obtained from NCBI Influenza Virus database (http://www.ncbi.nlm.nih.gov/genomes/FLU/FLU.html) and GISAID EpiFlu database (http://platform.gisaid.org), and the coding regions of these sequences were aligned using MAFFT v7.490. Before the analysis, a preliminary check using TempEst (v1.5.1) confirmed a significant temporal signal in our dataset, which was proved by linear regression of root-to-tip distance against sampling date (R2 = 0.8812, Correlation Coefficient = 0.9387). An uncorrelated lognormal relaxed molecular clock and a SRD06 nucleotide substitution model were implemented using the MCMC method run for 100 million generations and sampled every 1,0000 generations. Runs were assessed in Tracer v1.7.2 for sufficient convergence [effective sample size (ESS) > 200], and a maximum clade credibility (MCC) tree was generated in TreeAnnotator v1.10.4 after removing the first 10 % of runs as burn‐in. The obtained MCC tree was edited by FigTree v1.4.4

The GTR+F+G4 substitution model, which was selected using the Bayesian information criterion by ModelFinder in IQ-Tree v1.6.12, then the maximum likelihood (ML) trees of eight gene segments of our isolates were constructed with 1000 bootstraps. Genotype was confirmed in GenoFLU (https://github.com/USDA-VS/GenoFLU) (*3*).

**Figure S1-S8** are Phylogenic analysis of the NA and internal genes of H5N1 viruses by using Maximum likelihood method.

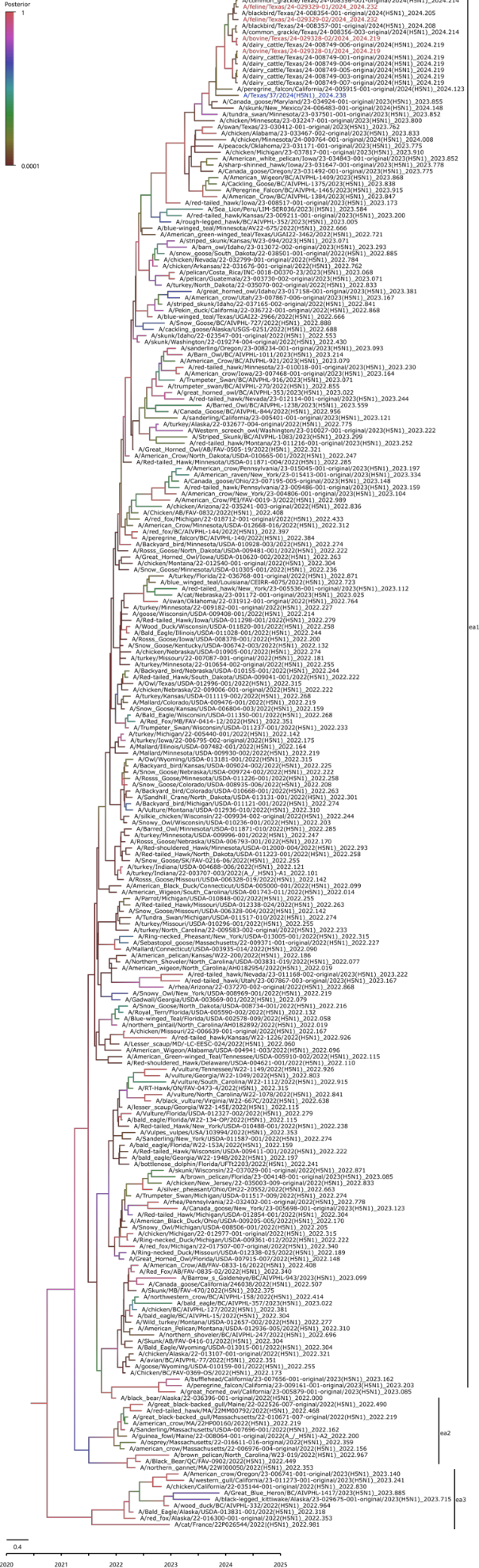

**Figure S1.** Phylogenic analysis of the HA segment of H5N1 viruses by using Maximum likelihood method. The H5N1 viruses isolated in this study are shown in red, human isolate is shown in blue and genotypic representatives in purple.

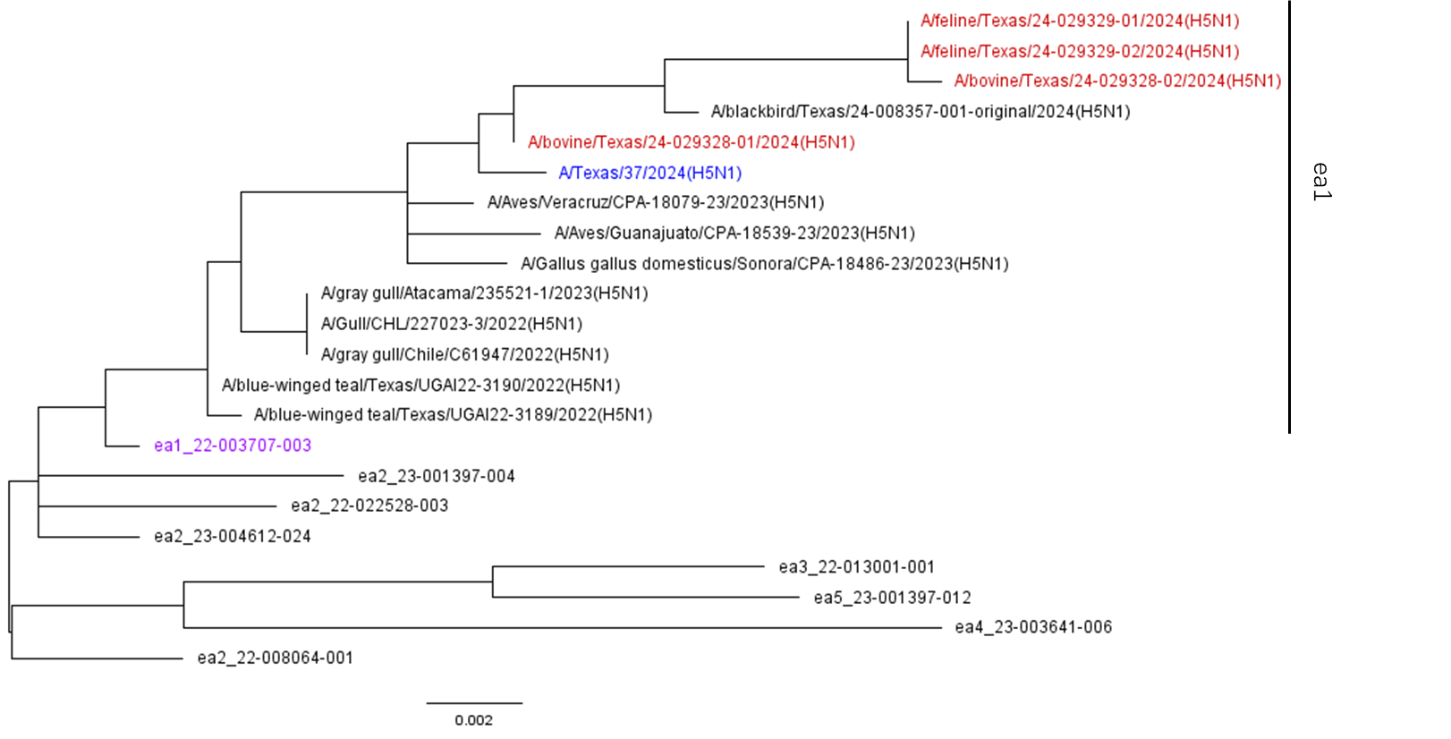

**Figure S2.** Phylogenic analysis of the NA segment of H5N1 viruses by using Maximum likelihood method. The H5N1 viruses isolated in this study are shown in red, human isolate is shown in blue and genotypic representatives in purple.

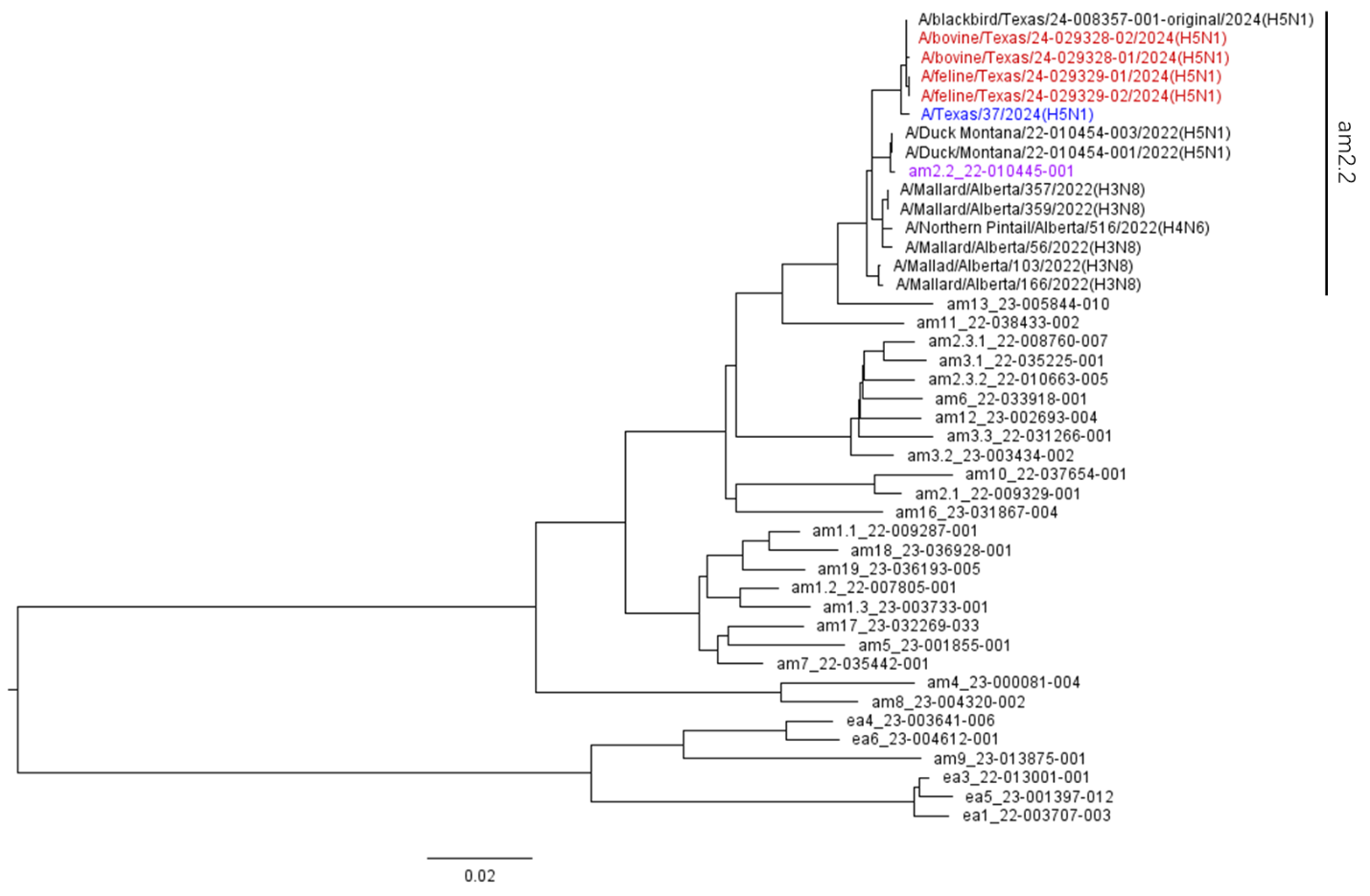

**Figure S3.** Phylogenic analysis of the PB2 segment of H5N1 viruses by using Maximum likelihood method. The H5N1 viruses isolated in this study are shown in red, human isolate is shown in blue and genotypic representatives in purple.

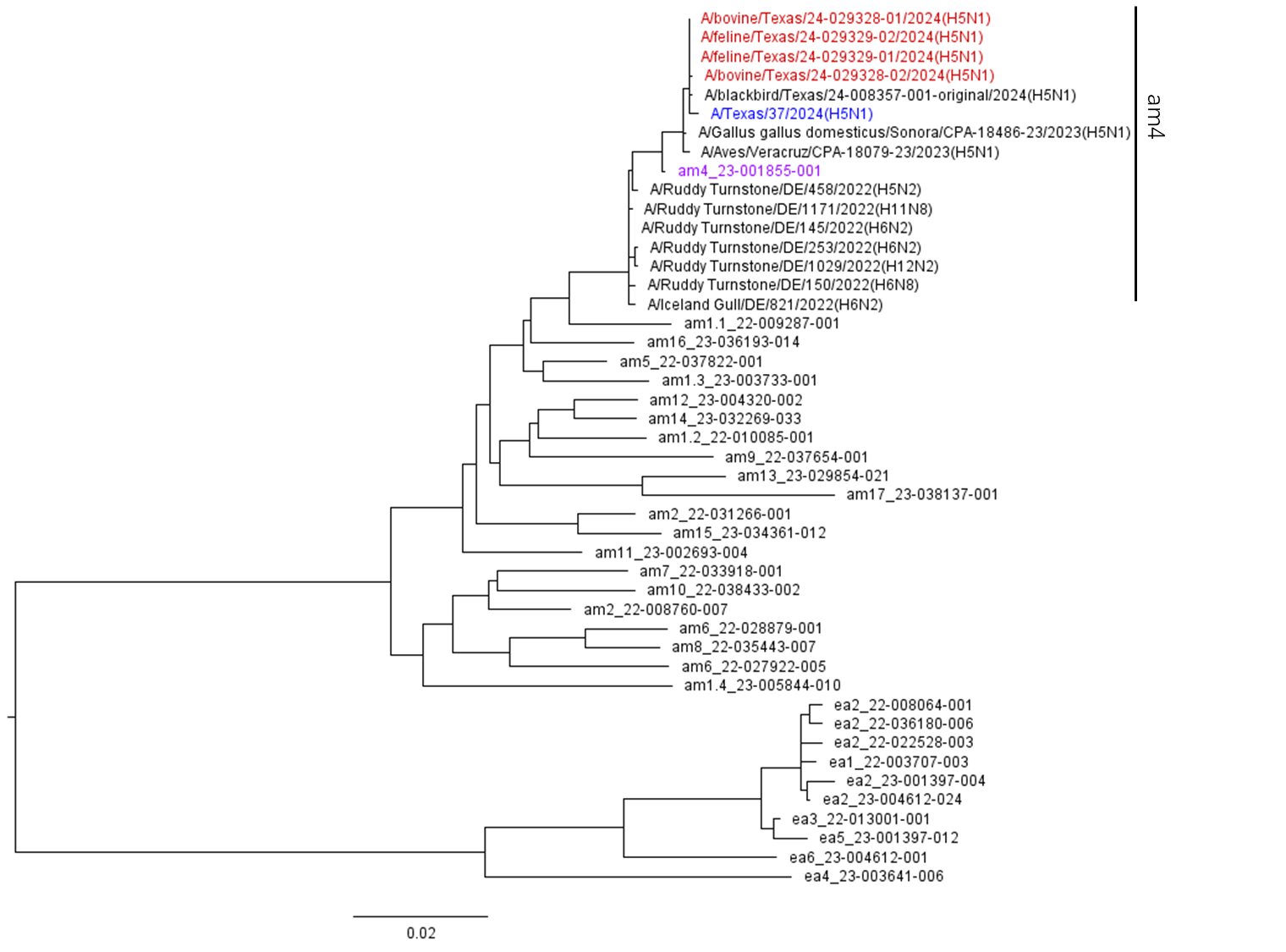

**Figure S4.** Phylogenic analysis of the PB1 segment of H5N1 viruses by using Maximum likelihood method. The H5N1 viruses isolated in this study are shown in red, human isolate is shown in blue and genotypic representatives in purple.

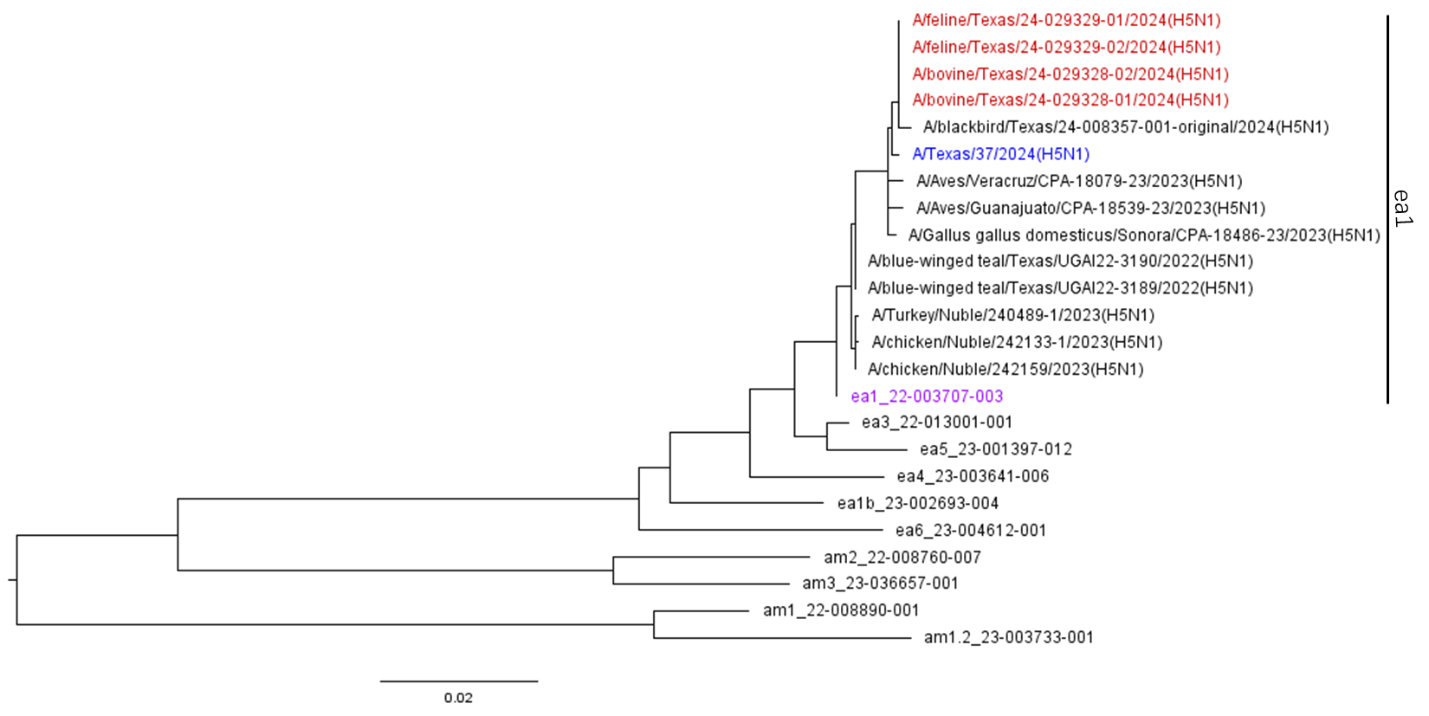

**Figure S5.** Phylogenic analysis of the PA segment of H5N1 viruses by using Maximum likelihood method. The H5N1 viruses isolated in this study are shown in red, human isolate is shown in blue and genotypic representatives in purple.

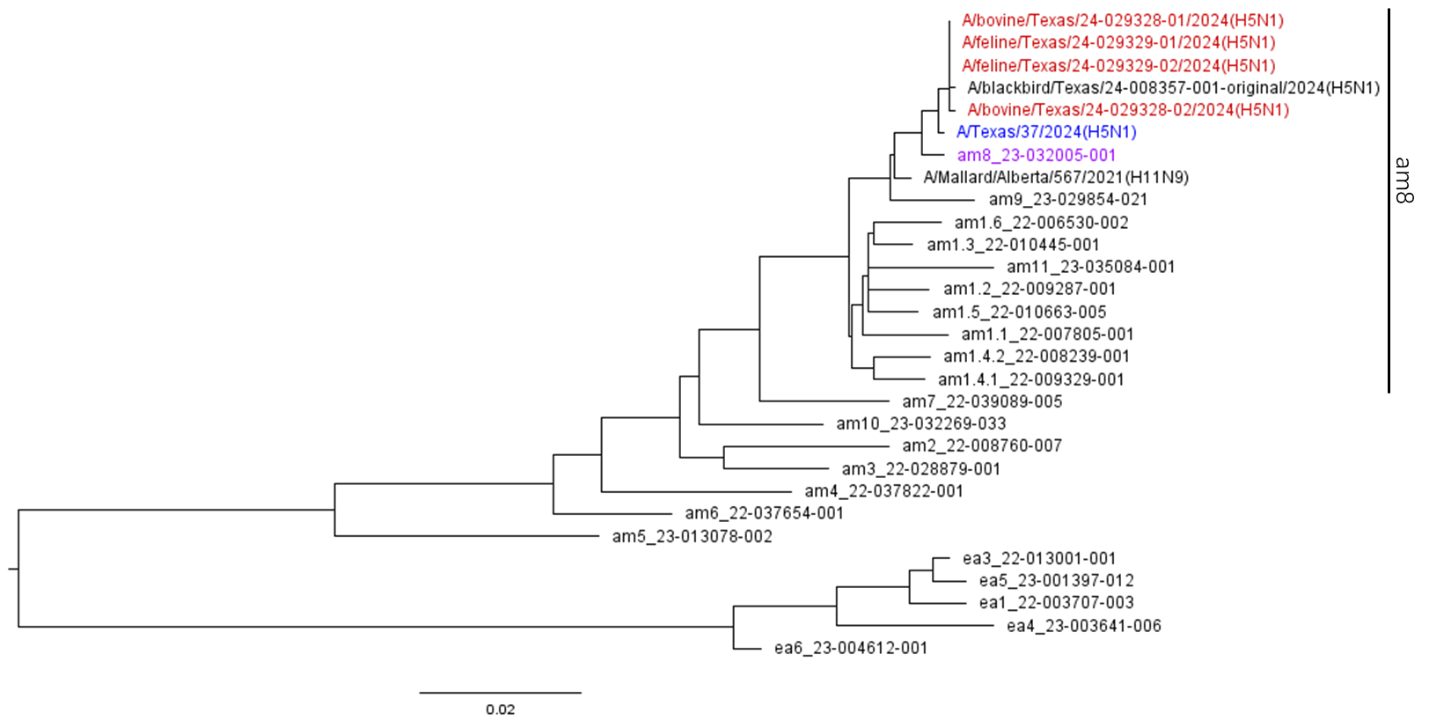

**Figure S6.** Phylogenic analysis of the NP segment of H5N1 viruses by using Maximum likelihood method. The H5N1 viruses isolated in this study are shown in red, human isolate is shown in blue and genotypic representatives in purple.

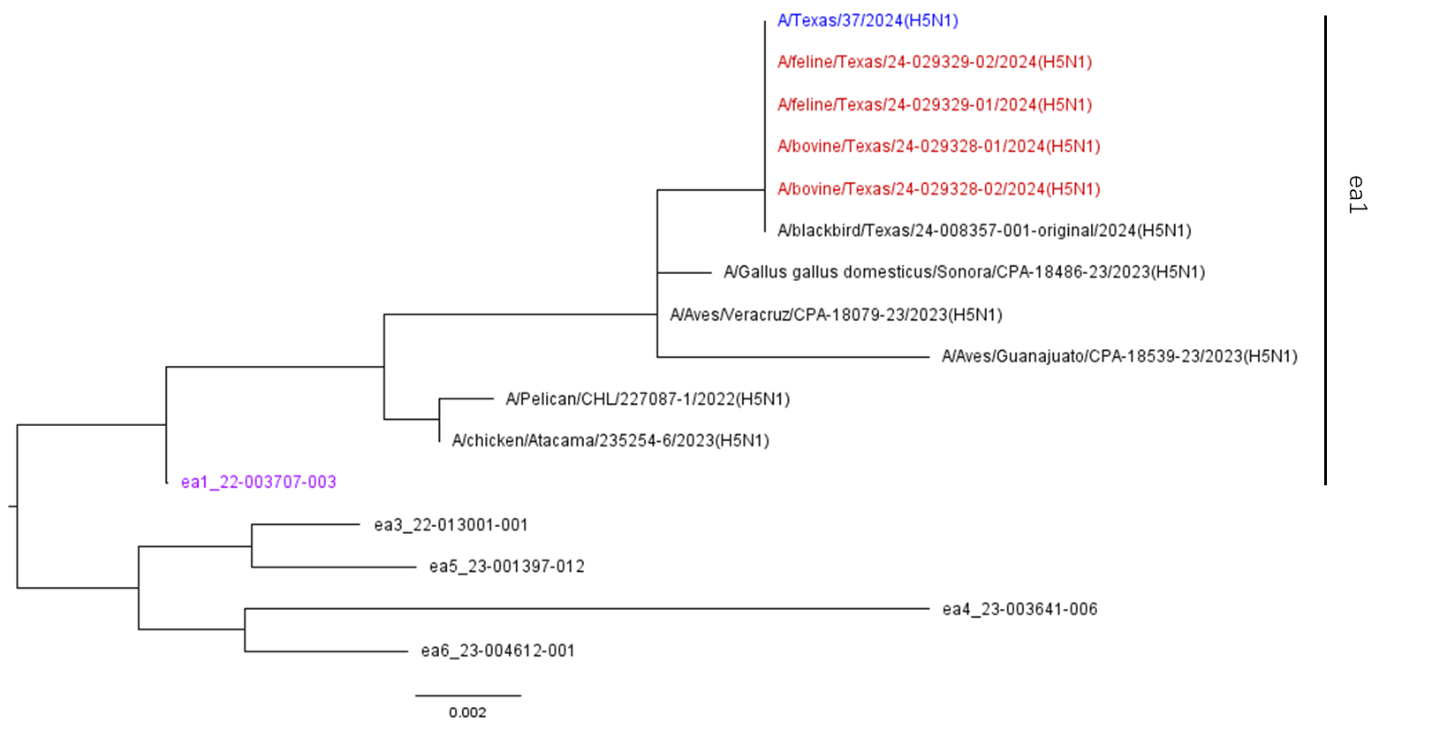

**Figure S7.** Phylogenic analysis of the M segment of H5N1 viruses by using Maximum likelihood method. The H5N1 viruses isolated in this study are shown in red, human isolate is shown in blue and genotypic representatives in purple.

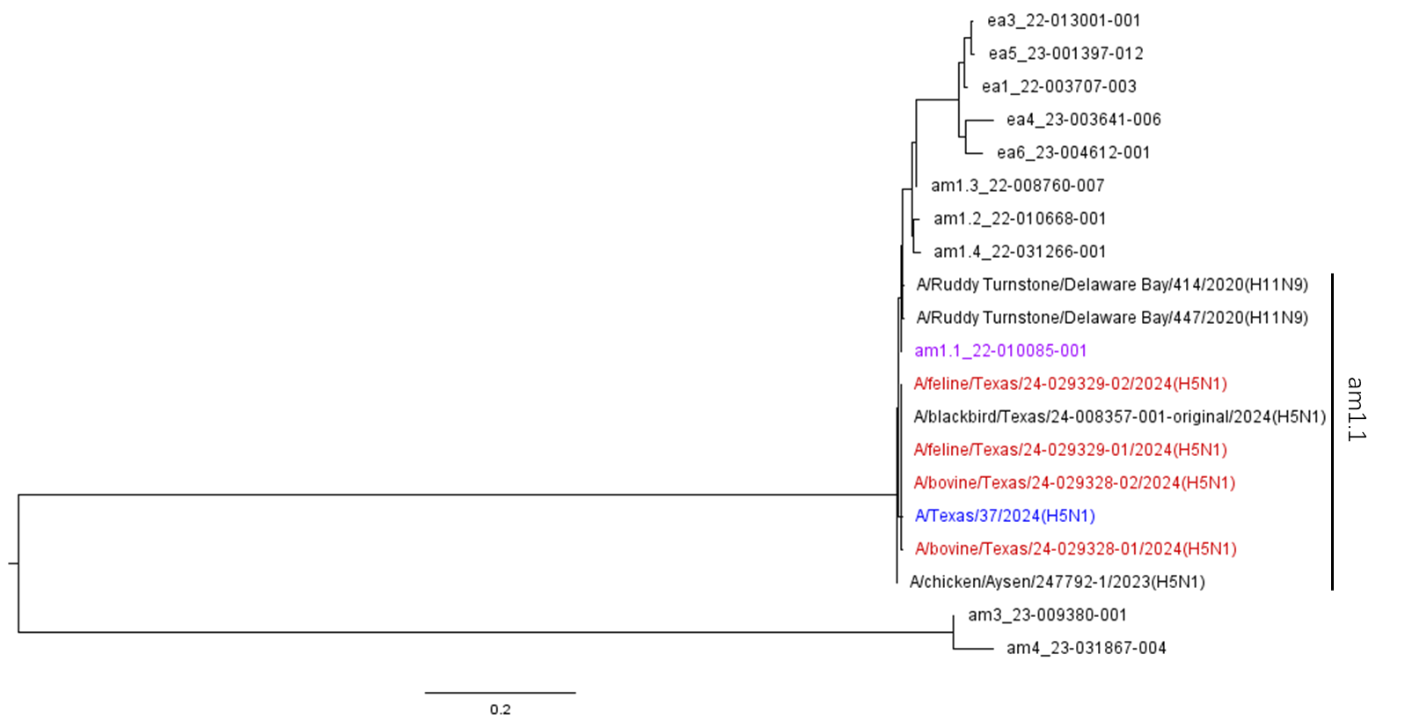

**Figure S8.** Phylogenic analysis of the NS segment of H5N1 viruses by using Maximum likelihood method. The H5N1 viruses isolated in this study are shown in red, human isolate is shown in blue and genotypic representatives in purple.

**Table S1.** Four HPAI A/H5N1 genome sequences from ISU-VDL in this study

| Virus Name | Abbr. | Species | Breed | Sample | Found | Sex | Age | Date | Location | Accession |
| --- | --- | --- | --- | --- | --- | --- | --- | --- | --- | --- |
| A/bovine/Texas/24-029328-01/2024 | Bovine-1 | Bovine | Jersey | Milk | Alive | Female | 3 Years | 3/20/2024 | Texas | SRR28464540 |
| A/bovine/Texas/24-029328-02/2024 | Bovine-2 | Bovine | Jersey | Milk | Alive | Female | 3 Years | 3/20/2024 | Texas | SRR28464539 |
| A/feline/Texas/24-029329-01/2024 | Feline-1 | Feline | Domestic Shorthair | Brain | Dead | N/A | 6 Months | 3/20/2024 | Texas | SRR28519493 |
| A/feline/Texas/24-029329-02/2024 | Feline-2 | Feline | Domestic Shorthair | Lung | Dead | N/A | 6 Months | 3/20/2024 | Texas | SRR28519492 |

**Table S2.** Fourteen HPAI A/H5N1 genome sequences submitted to GISAID by NVSLs that are included in this study

| Virus Name | Abbr. | PB2 | PB1 | PA | HA | NP | NA | MP | NS | Host | Species | Date | Location |
| --- | --- | --- | --- | --- | --- | --- | --- | --- | --- | --- | --- | --- | --- |
| A/dairy_cattle/Texas/24-008749-001-original/2024 | cattle-1 | EPI3158670 | EPI3158671 | EPI3158668 | EPI3158678 | EPI3158664 | EPI3158672 | EPI3158667 | EPI3158665 | Other mammals | Dairy cattle | 3/20/2024 | USA: Texas |
| A/dairy_cattle/Texas/24-008749-003-original/2024 | cattle-3 | EPI3158686 | EPI3158687 | EPI3158684 | EPI3158693 | EPI3158680 | EPI3158689 | EPI3158683 | EPI3158681 | Other mammals | Dairy cattle | 3/20/2024 | USA: Texas |
| A/dairy_cattle/Texas/24-008749-004-original/2024 | cattle-4 | EPI3158701 | EPI3158703 | EPI3158700 | EPI3158709 | EPI3158695 | EPI3158705 | EPI3158698 | EPI3158697 | Other mammals | Dairy cattle | 3/20/2024 | USA: Texas |
| A/dairy_cattle/Texas/24-008749-005-original/2024 | cattle-5 | EPI3158718 | EPI3158719 | EPI3158716 | EPI3158725 | EPI3158711 | EPI3158720 | EPI3158714 | EPI3158713 | Other mammals | Dairy cattle | 3/20/2024 | USA: Texas |
| A/dairy_cattle/Texas/24-008749-006-original/2024 | cattle-6 | EPI3158733 | EPI3158735 | EPI3158732 | EPI3158739 | EPI3158727 | EPI3158736 | EPI3158730 | EPI3158729 | Other mammals | Dairy cattle | 3/20/2024 | USA: Texas |
| A/dairy_cattle/Texas/24-008749-007-original/2024 | cattle-7 | EPI3158749 | EPI3158751 | EPI3158747 | EPI3158755 | EPI3158741 | EPI3158752 | EPI3158746 | EPI3158744 | Other mammals | Dairy cattle | 3/20/2024 | USA: Texas |
| A/skunk/New_Mexico/24-006483-001-original/2024 | skunk | EPI3158796 | EPI3158798 | EPI3158795 | EPI3158803 | EPI3158789 | EPI3158800 | EPI3158793 | EPI3158791 | Other mammals | Skunk | 2/23/2024 | USA: New Mexico |
| A/Texas/37/2024 | Human | EPI3171492 | EPI3171490 | EPI3171487 | EPI3171488 | EPI3171489 | EPI3171486 | EPI3171493 | EPI3171491 | Other mammals | Human | 3/28/2024 | USA: Texas |
| A/blackbird/Texas/24-008354-001-original/2024 | Blackbird-1 | EPI3158812 | EPI3158813 | EPI3158810 | EPI3158819 | EPI3158805 | EPI3158815 | EPI3158808 | EPI3158807 | Avian | Blackbird | 3/16/2024 | USA: Texas |
| A/blackbird/Texas/24-008357-001-original/2024 | Blackbird-1 | EPI3158860 | EPI3158861 | EPI3158858 | EPI3158865 | EPI3158853 | EPI3158863 | EPI3158856 | EPI3158855 | Avian | Blackbird | 3/16/2024 | USA: Texas |
| A/Canada_goose/Wyoming/24-003692-001-original/2024 | Goose | EPI3158764 | EPI3158766 | EPI3158763 | EPI3158771 | EPI3158757 | EPI3158768 | EPI3158761 | EPI3158759 | Avian | Canada goose | 1/25/2024 | USA: Wyoming |
| A/common_grackle/Texas/24-008356-001-original/2024 | Grackle-1 | EPI3158828 | EPI3158830 | EPI3158826 | EPI3158834 | EPI3158821 | EPI3158831 | EPI3158824 | EPI3158822 | Avian | Common grackle | 3/18/2024 | USA: Texas |
| A/common_grackle/Texas/24-008356-003-original/2024 | Grackle-2 | EPI3158844 | EPI3158846 | EPI3158842 | EPI3158851 | EPI3158836 | EPI3158847 | EPI3158841 | EPI3158839 | Avian | Common grackle | 3/18/2024 | USA: Texas |
| A/peregrine_falcon/California/24-005915-001-original/2024 | Falcon | EPI3158781 | EPI3158783 | EPI3158779 | EPI3158787 | EPI3158773 | EPI3158784 | EPI3158777 | EPI3158775 | Avian | Peregrine falcon | 2/14/2024 | USA: California |

**Table S3-S10** present the sequence identity matrices for each segment of the 18 HPAIV genomes. These matrices were computed using ssearch36 (version 36.3.8e), which facilitated the quantification of nucleotide sequence conservation across the viral isolates.

**Table S3.** Percentage identity of 18 HA segments

| **ID. for HA** | **Bovine-1** | **Bovine-2** | **Cattle-1** | **Cattle-3** | **Cattle-4** | **Cattle-5** | **Cattle-6** | **Cattle-7** | **Feline-1** | **Feline-2** | **Human** | **Skunk** | **Blackbird-1** | **Blackbird-2** | **Goose** | **Grackle-1** | **Grackle-2** | **Falcon** |
| --- | --- | --- | --- | --- | --- | --- | --- | --- | --- | --- | --- | --- | --- | --- | --- | --- | --- | --- |
| **Bovine-1** | **100** | **99.88** | **100** | **100** | **100** | **100** | **99.88** | **100** | **99.94** | **99.94** | **99.71** | **99.71** | **99.94** | **99.94** | **99.65** | **99.94** | **99.94** | **99.65** |
| **Bovine-2** | **99.88** | **100** | **99.88** | **99.88** | **99.88** | **99.88** | **100** | **99.88** | **99.94** | **99.94** | **99.71** | **99.71** | **99.94** | **99.94** | **99.65** | **99.94** | **99.94** | **99.65** |
| **Cattle-1** | **100** | **99.88** | **100** | **100** | **100** | **100** | **99.88** | **100** | **99.94** | **99.94** | **99.71** | **99.71** | **99.94** | **99.94** | **99.65** | **99.94** | **99.94** | **99.65** |
| **Cattle-3** | **100** | **99.88** | **100** | **100** | **100** | **100** | **99.88** | **100** | **99.94** | **99.94** | **99.71** | **99.71** | **99.94** | **99.94** | **99.65** | **99.94** | **99.94** | **99.65** |
| **Cattle-4** | **100** | **99.88** | **100** | **100** | **100** | **100** | **99.88** | **100** | **99.94** | **99.94** | **99.71** | **99.71** | **99.94** | **99.94** | **99.65** | **99.94** | **99.94** | **99.65** |
| **Cattle-5** | **100** | **99.88** | **100** | **100** | **100** | **100** | **99.88** | **100** | **99.94** | **99.94** | **99.71** | **99.71** | **99.94** | **99.94** | **99.65** | **99.94** | **99.94** | **99.65** |
| **Cattle-6** | **99.88** | **100** | **99.88** | **99.88** | **99.88** | **99.88** | **100** | **99.88** | **99.94** | **99.94** | **99.71** | **99.71** | **99.94** | **99.94** | **99.65** | **99.94** | **99.94** | **99.65** |
| **Cattle-7** | **100** | **99.88** | **100** | **100** | **100** | **100** | **99.88** | **100** | **99.94** | **99.94** | **99.71** | **99.71** | **99.94** | **99.94** | **99.65** | **99.94** | **99.94** | **99.65** |
| **Feline-1** | **99.94** | **99.94** | **99.94** | **99.94** | **99.94** | **99.94** | **99.94** | **99.94** | **100** | **100** | **99.77** | **99.77** | **100** | **100** | **99.71** | **100** | **100** | **99.71** |
| **Feline-2** | **99.94** | **99.94** | **99.94** | **99.94** | **99.94** | **99.94** | **99.94** | **99.94** | **100** | **100** | **99.77** | **99.77** | **100** | **100** | **99.71** | **100** | **100** | **99.71** |
| **Human** | **99.71** | **99.71** | **99.71** | **99.71** | **99.71** | **99.71** | **99.71** | **99.71** | **99.77** | **99.77** | **100** | **99.77** | **99.77** | **99.77** | **99.71** | **99.77** | **99.77** | **99.71** |
| **Skunk** | **99.71** | **99.71** | **99.71** | **99.71** | **99.71** | **99.71** | **99.71** | **99.71** | **99.77** | **99.77** | **99.77** | **100** | **99.77** | **99.77** | **99.71** | **99.77** | **99.77** | **99.71** |
| **Blackbird-1** | **99.94** | **99.94** | **99.94** | **99.94** | **99.94** | **99.94** | **99.94** | **99.94** | **100** | **100** | **99.77** | **99.77** | **100** | **100** | **99.71** | **100** | **100** | **99.71** |
| **Blackbird-2** | **99.94** | **99.94** | **99.94** | **99.94** | **99.94** | **99.94** | **99.94** | **99.94** | **100** | **100** | **99.77** | **99.77** | **100** | **100** | **99.71** | **100** | **100** | **99.71** |
| **Goose** | **99.65** | **99.65** | **99.65** | **99.65** | **99.65** | **99.65** | **99.65** | **99.65** | **99.71** | **99.71** | **99.71** | **99.71** | **99.71** | **99.71** | **100** | **99.71** | **99.71** | **99.65** |
| **Grackle-1** | **99.94** | **99.94** | **99.94** | **99.94** | **99.94** | **99.94** | **99.94** | **99.94** | **100** | **100** | **99.77** | **99.77** | **100** | **100** | **99.71** | **100** | **100** | **99.71** |
| **Grackle-2** | **99.94** | **99.94** | **99.94** | **99.94** | **99.94** | **99.94** | **99.94** | **99.94** | **100** | **100** | **99.77** | **99.77** | **100** | **100** | **99.71** | **100** | **100** | **99.71** |
| **Falcon** | **99.65** | **99.65** | **99.65** | **99.65** | **99.65** | **99.65** | **99.65** | **99.65** | **99.71** | **99.71** | **99.71** | **99.71** | **99.71** | **99.71** | **99.65** | **99.71** | **99.71** | **100** |

**Table S4.** Percentage identity of 18 NA segments

| **ID. for NA** | **Bovine-1** | **Bovine-2** | **Cattle-1** | **Cattle-3** | **Cattle-4** | **Cattle-5** | **Cattle-6** | **Cattle-7** | **Feline-1** | **Feline-2** | **Human** | **Skunk** | **Blackbird-1** | **Blackbird-2** | **Goose** | **Grackle-1** | **Grackle-2** | **Falcon** |
| --- | --- | --- | --- | --- | --- | --- | --- | --- | --- | --- | --- | --- | --- | --- | --- | --- | --- | --- |
| **Bovine-1** | **100** | **99.93** | **100** | **100** | **100** | **100** | **99.93** | **100** | **100** | **100** | **99.79** | **99.65** | **100** | **99.93** | **99.79** | **99.86** | **99.79** | **99.5** |
| **Bovine-2** | **99.93** | **100** | **99.93** | **99.93** | **99.93** | **99.93** | **100** | **99.93** | **99.93** | **99.93** | **99.72** | **99.57** | **99.93** | **99.86** | **99.72** | **99.79** | **99.72** | **99.43** |
| **Cattle-1** | **100** | **99.93** | **100** | **100** | **100** | **100** | **99.93** | **100** | **100** | **100** | **99.79** | **99.65** | **100** | **99.93** | **99.79** | **99.86** | **99.79** | **99.5** |
| **Cattle-3** | **100** | **99.93** | **100** | **100** | **100** | **100** | **99.93** | **100** | **100** | **100** | **99.79** | **99.65** | **100** | **99.93** | **99.79** | **99.86** | **99.79** | **99.5** |
| **Cattle-4** | **100** | **99.93** | **100** | **100** | **100** | **100** | **99.93** | **100** | **100** | **100** | **99.79** | **99.65** | **100** | **99.93** | **99.79** | **99.86** | **99.79** | **99.5** |
| **Cattle-5** | **100** | **99.93** | **100** | **100** | **100** | **100** | **99.93** | **100** | **100** | **100** | **99.79** | **99.65** | **100** | **99.93** | **99.79** | **99.86** | **99.79** | **99.5** |
| **Cattle-6** | **99.93** | **100** | **99.93** | **99.93** | **99.93** | **99.93** | **100** | **99.93** | **99.93** | **99.93** | **99.72** | **99.57** | **99.93** | **99.86** | **99.72** | **99.79** | **99.72** | **99.43** |
| **Cattle-7** | **100** | **99.93** | **100** | **100** | **100** | **100** | **99.93** | **100** | **100** | **100** | **99.79** | **99.65** | **100** | **99.93** | **99.79** | **99.86** | **99.79** | **99.5** |
| **Feline-1** | **100** | **99.93** | **100** | **100** | **100** | **100** | **99.93** | **100** | **100** | **100** | **99.79** | **99.65** | **100** | **99.93** | **99.79** | **99.86** | **99.79** | **99.5** |
| **Feline-2** | **100** | **99.93** | **100** | **100** | **100** | **100** | **99.93** | **100** | **100** | **100** | **99.79** | **99.65** | **100** | **99.93** | **99.79** | **99.86** | **99.79** | **99.5** |
| **Human** | **99.79** | **99.72** | **99.79** | **99.79** | **99.79** | **99.79** | **99.72** | **99.79** | **99.79** | **99.79** | **100** | **99.57** | **99.79** | **99.72** | **99.72** | **99.79** | **99.72** | **99.43** |
| **Skunk** | **99.65** | **99.57** | **99.65** | **99.65** | **99.65** | **99.65** | **99.57** | **99.65** | **99.65** | **99.65** | **99.57** | **100** | **99.65** | **99.57** | **99.72** | **99.65** | **99.57** | **99.43** |
| **Blackbird-1** | **100** | **99.93** | **100** | **100** | **100** | **100** | **99.93** | **100** | **100** | **100** | **99.79** | **99.65** | **100** | **99.93** | **99.79** | **99.86** | **99.79** | **99.5** |
| **Blackbird-2** | **99.93** | **99.86** | **99.93** | **99.93** | **99.93** | **99.93** | **99.86** | **99.93** | **99.93** | **99.93** | **99.72** | **99.57** | **99.93** | **100** | **99.72** | **99.79** | **99.72** | **99.43** |
| **Goose** | **99.79** | **99.72** | **99.79** | **99.79** | **99.79** | **99.79** | **99.72** | **99.79** | **99.79** | **99.79** | **99.72** | **99.72** | **99.79** | **99.72** | **100** | **99.79** | **99.72** | **99.57** |
| **Grackle-1** | **99.86** | **99.79** | **99.86** | **99.86** | **99.86** | **99.86** | **99.79** | **99.86** | **99.86** | **99.86** | **99.79** | **99.65** | **99.86** | **99.79** | **99.79** | **100** | **99.93** | **99.5** |
| **Grackle-2** | **99.79** | **99.72** | **99.79** | **99.79** | **99.79** | **99.79** | **99.72** | **99.79** | **99.79** | **99.79** | **99.72** | **99.57** | **99.79** | **99.72** | **99.72** | **99.93** | **100** | **99.43** |
| **Falcon** | **99.5** | **99.43** | **99.5** | **99.5** | **99.5** | **99.5** | **99.43** | **99.5** | **99.5** | **99.5** | **99.43** | **99.43** | **99.5** | **99.43** | **99.57** | **99.5** | **99.43** | **100** |

**Table S5.** Percentage identity of 18 NS segments

| **dID. for NS** | **Bovine-1** | **Bovine-2** | **Cattle-1** | **Cattle-3** | **Cattle-4** | **Cattle-5** | **Cattle-6** | **Cattle-7** | **Feline-1** | **Feline-2** | **Human** | **Skunk** | **Blackbird-1** | **Blackbird-2** | **Goose** | **Grackle-1** | **Grackle-2** | **Falcon** |
| --- | --- | --- | --- | --- | --- | --- | --- | --- | --- | --- | --- | --- | --- | --- | --- | --- | --- | --- |
| **Bovine-1** | **100** | **99.88** | **100** | **100** | **100** | **100** | **99.88** | **100** | **99.88** | **99.88** | **99.64** | **99.76** | **99.76** | **99.88** | **99.64** | **99.76** | **99.76** | **99.76** |
| **Bovine-2** | **99.88** | **100** | **99.88** | **99.88** | **99.88** | **99.88** | **100** | **99.88** | **100** | **100** | **99.76** | **99.88** | **99.88** | **100** | **99.76** | **99.88** | **99.88** | **99.88** |
| **Cattle-1** | **100** | **99.88** | **100** | **100** | **100** | **100** | **99.88** | **100** | **99.88** | **99.88** | **99.64** | **99.76** | **99.76** | **99.88** | **99.64** | **99.76** | **99.76** | **99.76** |
| **Cattle-3** | **100** | **99.88** | **100** | **100** | **100** | **100** | **99.88** | **100** | **99.88** | **99.88** | **99.64** | **99.76** | **99.76** | **99.88** | **99.64** | **99.76** | **99.76** | **99.76** |
| **Cattle-4** | **100** | **99.88** | **100** | **100** | **100** | **100** | **99.88** | **100** | **99.88** | **99.88** | **99.64** | **99.76** | **99.76** | **99.88** | **99.64** | **99.76** | **99.76** | **99.76** |
| **Cattle-5** | **100** | **99.88** | **100** | **100** | **100** | **100** | **99.88** | **100** | **99.88** | **99.88** | **99.64** | **99.76** | **99.76** | **99.88** | **99.64** | **99.76** | **99.76** | **99.76** |
| **Cattle-6** | **99.88** | **100** | **99.88** | **99.88** | **99.88** | **99.88** | **100** | **99.88** | **100** | **100** | **99.76** | **99.88** | **99.88** | **100** | **99.76** | **99.88** | **99.88** | **99.88** |
| **Cattle-7** | **100** | **99.88** | **100** | **100** | **100** | **100** | **99.88** | **100** | **99.88** | **99.88** | **99.64** | **99.76** | **99.76** | **99.88** | **99.64** | **99.76** | **99.76** | **99.76** |
| **Feline-1** | **99.88** | **100** | **99.88** | **99.88** | **99.88** | **99.88** | **100** | **99.88** | **100** | **100** | **99.76** | **99.88** | **99.88** | **100** | **99.76** | **99.88** | **99.88** | **99.88** |
| **Feline-2** | **99.88** | **100** | **99.88** | **99.88** | **99.88** | **99.88** | **100** | **99.88** | **100** | **100** | **99.76** | **99.88** | **99.88** | **100** | **99.76** | **99.88** | **99.88** | **99.88** |
| **Human** | **99.64** | **99.76** | **99.64** | **99.64** | **99.64** | **99.64** | **99.76** | **99.64** | **99.76** | **99.76** | **100** | **99.64** | **99.64** | **99.76** | **99.52** | **99.64** | **99.64** | **99.64** |
| **Skunk** | **99.76** | **99.88** | **99.76** | **99.76** | **99.76** | **99.76** | **99.88** | **99.76** | **99.88** | **99.88** | **99.64** | **100** | **99.76** | **99.88** | **99.88** | **99.76** | **99.76** | **100** |
| **Blackbird-1** | **99.76** | **99.88** | **99.76** | **99.76** | **99.76** | **99.76** | **99.88** | **99.76** | **99.88** | **99.88** | **99.64** | **99.76** | **100** | **99.88** | **99.64** | **99.76** | **99.76** | **99.76** |
| **Blackbird-2** | **99.88** | **100** | **99.88** | **99.88** | **99.88** | **99.88** | **100** | **99.88** | **100** | **100** | **99.76** | **99.88** | **99.88** | **100** | **99.76** | **99.88** | **99.88** | **99.88** |
| **Goose** | **99.64** | **99.76** | **99.64** | **99.64** | **99.64** | **99.64** | **99.76** | **99.64** | **99.76** | **99.76** | **99.52** | **99.88** | **99.64** | **99.76** | **100** | **99.64** | **99.64** | **99.88** |
| **Grackle-1** | **99.76** | **99.88** | **99.76** | **99.76** | **99.76** | **99.76** | **99.88** | **99.76** | **99.88** | **99.88** | **99.64** | **99.76** | **99.76** | **99.88** | **99.64** | **100** | **100** | **99.76** |
| **Grackle-2** | **99.76** | **99.88** | **99.76** | **99.76** | **99.76** | **99.76** | **99.88** | **99.76** | **99.88** | **99.88** | **99.64** | **99.76** | **99.76** | **99.88** | **99.64** | **100** | **100** | **99.76** |
| **Falcon** | **99.76** | **99.88** | **99.76** | **99.76** | **99.76** | **99.76** | **99.88** | **99.76** | **99.88** | **99.88** | **99.64** | **100** | **99.76** | **99.88** | **99.88** | **99.76** | **99.76** | **100** |

**Table S6.** Percentage identity of 18 M segments

| **ID. for M** | **Bovine-1** | **Bovine-2** | **Cattle-1** | **Cattle-3** | **Cattle-4** | **Cattle-5** | **Cattle-6** | **Cattle-7** | **Feline-1** | **Feline-2** | **Human** | **Skunk** | **Blackbird-1** | **Blackbird-2** | **Goose** | **Grackle-1** | **Grackle-2** | **Falcon** |
| --- | --- | --- | --- | --- | --- | --- | --- | --- | --- | --- | --- | --- | --- | --- | --- | --- | --- | --- |
| **Bovine-1** | **100** | **100** | **100** | **100** | **100** | **100** | **100** | **100** | **100** | **100** | **100** | **99.8** | **100** | **100** | **99.8** | **100** | **100** | **99.69** |
| **Bovine-2** | **100** | **100** | **100** | **100** | **100** | **100** | **100** | **100** | **100** | **100** | **100** | **99.8** | **100** | **100** | **99.8** | **100** | **100** | **99.69** |
| **Cattle-1** | **100** | **100** | **100** | **100** | **100** | **100** | **100** | **100** | **100** | **100** | **100** | **99.8** | **100** | **100** | **99.8** | **100** | **100** | **99.69** |
| **Cattle-3** | **100** | **100** | **100** | **100** | **100** | **100** | **100** | **100** | **100** | **100** | **100** | **99.8** | **100** | **100** | **99.8** | **100** | **100** | **99.69** |
| **Cattle-4** | **100** | **100** | **100** | **100** | **100** | **100** | **100** | **100** | **100** | **100** | **100** | **99.8** | **100** | **100** | **99.8** | **100** | **100** | **99.69** |
| **Cattle-5** | **100** | **100** | **100** | **100** | **100** | **100** | **100** | **100** | **100** | **100** | **100** | **99.8** | **100** | **100** | **99.8** | **100** | **100** | **99.69** |
| **Cattle-6** | **100** | **100** | **100** | **100** | **100** | **100** | **100** | **100** | **100** | **100** | **100** | **99.8** | **100** | **100** | **99.8** | **100** | **100** | **99.69** |
| **Cattle-7** | **100** | **100** | **100** | **100** | **100** | **100** | **100** | **100** | **100** | **100** | **100** | **99.8** | **100** | **100** | **99.8** | **100** | **100** | **99.69** |
| **Feline-1** | **100** | **100** | **100** | **100** | **100** | **100** | **100** | **100** | **100** | **100** | **100** | **99.8** | **100** | **100** | **99.8** | **100** | **100** | **99.69** |
| **Feline-2** | **100** | **100** | **100** | **100** | **100** | **100** | **100** | **100** | **100** | **100** | **100** | **99.8** | **100** | **100** | **99.8** | **100** | **100** | **99.69** |
| **Human** | **100** | **100** | **100** | **100** | **100** | **100** | **100** | **100** | **100** | **100** | **100** | **99.8** | **100** | **100** | **99.8** | **100** | **100** | **99.69** |
| **Skunk** | **99.8** | **99.8** | **99.8** | **99.8** | **99.8** | **99.8** | **99.8** | **99.8** | **99.8** | **99.8** | **99.8** | **100** | **99.8** | **99.8** | **100** | **99.8** | **99.8** | **99.9** |
| **Blackbird-1** | **100** | **100** | **100** | **100** | **100** | **100** | **100** | **100** | **100** | **100** | **100** | **99.8** | **100** | **100** | **99.8** | **100** | **100** | **99.69** |
| **Blackbird-2** | **100** | **100** | **100** | **100** | **100** | **100** | **100** | **100** | **100** | **100** | **100** | **99.8** | **100** | **100** | **99.8** | **100** | **100** | **99.69** |
| **Goose** | **99.8** | **99.8** | **99.8** | **99.8** | **99.8** | **99.8** | **99.8** | **99.8** | **99.8** | **99.8** | **99.8** | **100** | **99.8** | **99.8** | **100** | **99.8** | **99.8** | **99.9** |
| **Grackle-1** | **100** | **100** | **100** | **100** | **100** | **100** | **100** | **100** | **100** | **100** | **100** | **99.8** | **100** | **100** | **99.8** | **100** | **100** | **99.69** |
| **Grackle-2** | **100** | **100** | **100** | **100** | **100** | **100** | **100** | **100** | **100** | **100** | **100** | **99.8** | **100** | **100** | **99.8** | **100** | **100** | **99.69** |
| **Falcon** | **99.69** | **99.69** | **99.69** | **99.69** | **99.69** | **99.69** | **99.69** | **99.69** | **99.69** | **99.69** | **99.69** | **99.9** | **99.69** | **99.69** | **99.9** | **99.69** | **99.69** | **100** |

**Table S7.** Percentage identity of 18 NP segments

| **ID. for NP** | **Bovine-1** | **Bovine-2** | **Cattle-1** | **Cattle-3** | **Cattle-4** | **Cattle-5** | **Cattle-6** | **Cattle-7** | **Feline-1** | **Feline-2** | **Human** | **Skunk** | **Blackbird-1** | **Blackbird-2** | **Goose** | **Grackle-1** | **Grackle-2** | **Falcon** |
| --- | --- | --- | --- | --- | --- | --- | --- | --- | --- | --- | --- | --- | --- | --- | --- | --- | --- | --- |
| **Bovine-1** | **100** | **99.93** | **100** | **100** | **100** | **100** | **99.93** | **100** | **100** | **100** | **99.8** | **99.73** | **99.93** | **99.93** | **99.67** | **99.93** | **99.87** | **99.73** |
| **Bovine-2** | **99.93** | **100** | **99.93** | **99.93** | **99.93** | **99.93** | **100** | **99.93** | **99.93** | **99.93** | **99.73** | **99.67** | **99.87** | **99.87** | **99.6** | **99.87** | **99.8** | **99.67** |
| **Cattle-1** | **100** | **99.93** | **100** | **100** | **100** | **100** | **99.93** | **100** | **100** | **100** | **99.8** | **99.73** | **99.93** | **99.93** | **99.67** | **99.93** | **99.87** | **99.73** |
| **Cattle-3** | **100** | **99.93** | **100** | **100** | **100** | **100** | **99.93** | **100** | **100** | **100** | **99.8** | **99.73** | **99.93** | **99.93** | **99.67** | **99.93** | **99.87** | **99.73** |
| **Cattle-4** | **100** | **99.93** | **100** | **100** | **100** | **100** | **99.93** | **100** | **100** | **100** | **99.8** | **99.73** | **99.93** | **99.93** | **99.67** | **99.93** | **99.87** | **99.73** |
| **Cattle-5** | **100** | **99.93** | **100** | **100** | **100** | **100** | **99.93** | **100** | **100** | **100** | **99.8** | **99.73** | **99.93** | **99.93** | **99.67** | **99.93** | **99.87** | **99.73** |
| **Cattle-6** | **99.93** | **100** | **99.93** | **99.93** | **99.93** | **99.93** | **100** | **99.93** | **99.93** | **99.93** | **99.73** | **99.67** | **99.87** | **99.87** | **99.6** | **99.87** | **99.8** | **99.67** |
| **Cattle-7** | **100** | **99.93** | **100** | **100** | **100** | **100** | **99.93** | **100** | **100** | **100** | **99.8** | **99.73** | **99.93** | **99.93** | **99.67** | **99.93** | **99.87** | **99.73** |
| **Feline-1** | **100** | **99.93** | **100** | **100** | **100** | **100** | **99.93** | **100** | **100** | **100** | **99.8** | **99.73** | **99.93** | **99.93** | **99.67** | **99.93** | **99.87** | **99.73** |
| **Feline-2** | **100** | **99.93** | **100** | **100** | **100** | **100** | **99.93** | **100** | **100** | **100** | **99.8** | **99.73** | **99.93** | **99.93** | **99.67** | **99.93** | **99.87** | **99.73** |
| **Human** | **99.8** | **99.73** | **99.8** | **99.8** | **99.8** | **99.8** | **99.73** | **99.8** | **99.8** | **99.8** | **100** | **99.8** | **99.73** | **99.73** | **99.73** | **99.73** | **99.67** | **99.93** |
| **Skunk** | **99.73** | **99.67** | **99.73** | **99.73** | **99.73** | **99.73** | **99.67** | **99.73** | **99.73** | **99.73** | **99.8** | **100** | **99.67** | **99.67** | **99.8** | **99.67** | **99.6** | **99.87** |
| **Blackbird-1** | **99.93** | **99.87** | **99.93** | **99.93** | **99.93** | **99.93** | **99.87** | **99.93** | **99.93** | **99.93** | **99.73** | **99.67** | **100** | **99.87** | **99.6** | **99.87** | **99.8** | **99.67** |
| **Blackbird-2** | **99.93** | **99.87** | **99.93** | **99.93** | **99.93** | **99.93** | **99.87** | **99.93** | **99.93** | **99.93** | **99.73** | **99.67** | **99.87** | **100** | **99.6** | **99.87** | **99.8** | **99.67** |
| **Goose** | **99.67** | **99.6** | **99.67** | **99.67** | **99.67** | **99.67** | **99.6** | **99.67** | **99.67** | **99.67** | **99.73** | **99.8** | **99.6** | **99.6** | **100** | **99.6** | **99.53** | **99.8** |
| **Grackle-1** | **99.93** | **99.87** | **99.93** | **99.93** | **99.93** | **99.93** | **99.87** | **99.93** | **99.93** | **99.93** | **99.73** | **99.67** | **99.87** | **99.87** | **99.6** | **100** | **99.93** | **99.67** |
| **Grackle-2** | **99.87** | **99.8** | **99.87** | **99.87** | **99.87** | **99.87** | **99.8** | **99.87** | **99.87** | **99.87** | **99.67** | **99.6** | **99.8** | **99.8** | **99.53** | **99.93** | **100** | **99.6** |
| **Falcon** | **99.73** | **99.67** | **99.73** | **99.73** | **99.73** | **99.73** | **99.67** | **99.73** | **99.73** | **99.73** | **99.93** | **99.87** | **99.67** | **99.67** | **99.8** | **99.67** | **99.6** | **100** |

**Table S8.** Percentage identity of 18 PA segments

| **ID. for PA** | **Bovine-1** | **Bovine-2** | **Cattle-1** | **Cattle-3** | **Cattle-4** | **Cattle-5** | **Cattle-6** | **Cattle-7** | **Feline-1** | **Feline-2** | **Human** | **Skunk** | **Blackbird-1** | **Blackbird-2** | **Goose** | **Grackle-1** | **Grackle-2** | **Falcon** |
| --- | --- | --- | --- | --- | --- | --- | --- | --- | --- | --- | --- | --- | --- | --- | --- | --- | --- | --- |
| **Bovine-1** | **100** | **100** | **100** | **100** | **100** | **100** | **100** | **100** | **100** | **100** | **99.81** | **99.81** | **99.95** | **99.86** | **99.72** | **100** | **99.95** | **99.81** |
| **Bovine-2** | **100** | **100** | **100** | **100** | **100** | **100** | **100** | **100** | **100** | **100** | **99.81** | **99.81** | **99.95** | **99.86** | **99.72** | **100** | **99.95** | **99.81** |
| **Cattle-1** | **100** | **100** | **100** | **100** | **100** | **100** | **100** | **100** | **100** | **100** | **99.81** | **99.81** | **99.95** | **99.86** | **99.72** | **100** | **99.95** | **99.81** |
| **Cattle-3** | **100** | **100** | **100** | **100** | **100** | **100** | **100** | **100** | **100** | **100** | **99.81** | **99.81** | **99.95** | **99.86** | **99.72** | **100** | **99.95** | **99.81** |
| **Cattle-4** | **100** | **100** | **100** | **100** | **100** | **100** | **100** | **100** | **100** | **100** | **99.81** | **99.81** | **99.95** | **99.86** | **99.72** | **100** | **99.95** | **99.81** |
| **Cattle-5** | **100** | **100** | **100** | **100** | **100** | **100** | **100** | **100** | **100** | **100** | **99.81** | **99.81** | **99.95** | **99.86** | **99.72** | **100** | **99.95** | **99.81** |
| **Cattle-6** | **100** | **100** | **100** | **100** | **100** | **100** | **100** | **100** | **100** | **100** | **99.81** | **99.81** | **99.95** | **99.86** | **99.72** | **100** | **99.95** | **99.81** |
| **Cattle-7** | **100** | **100** | **100** | **100** | **100** | **100** | **100** | **100** | **100** | **100** | **99.81** | **99.81** | **99.95** | **99.86** | **99.72** | **100** | **99.95** | **99.81** |
| **Feline-1** | **100** | **100** | **100** | **100** | **100** | **100** | **100** | **100** | **100** | **100** | **99.81** | **99.81** | **99.95** | **99.86** | **99.72** | **100** | **99.95** | **99.81** |
| **Feline-2** | **100** | **100** | **100** | **100** | **100** | **100** | **100** | **100** | **100** | **100** | **99.81** | **99.81** | **99.95** | **99.86** | **99.72** | **100** | **99.95** | **99.81** |
| **Human** | **99.81** | **99.81** | **99.81** | **99.81** | **99.81** | **99.81** | **99.81** | **99.81** | **99.81** | **99.81** | **100** | **99.81** | **99.77** | **99.67** | **99.72** | **99.81** | **99.77** | **99.81** |
| **Skunk** | **99.81** | **99.81** | **99.81** | **99.81** | **99.81** | **99.81** | **99.81** | **99.81** | **99.81** | **99.81** | **99.81** | **100** | **99.77** | **99.67** | **99.72** | **99.81** | **99.77** | **99.81** |
| **Blackbird-1** | **99.95** | **99.95** | **99.95** | **99.95** | **99.95** | **99.95** | **99.95** | **99.95** | **99.95** | **99.95** | **99.77** | **99.77** | **100** | **99.81** | **99.67** | **99.95** | **99.91** | **99.77** |
| **Blackbird-2** | **99.86** | **99.86** | **99.86** | **99.86** | **99.86** | **99.86** | **99.86** | **99.86** | **99.86** | **99.86** | **99.67** | **99.67** | **99.81** | **100** | **99.58** | **99.86** | **99.81** | **99.67** |
| **Goose** | **99.72** | **99.72** | **99.72** | **99.72** | **99.72** | **99.72** | **99.72** | **99.72** | **99.72** | **99.72** | **99.72** | **99.72** | **99.67** | **99.58** | **100** | **99.72** | **99.67** | **99.72** |
| **Grackle-1** | **100** | **100** | **100** | **100** | **100** | **100** | **100** | **100** | **100** | **100** | **99.81** | **99.81** | **99.95** | **99.86** | **99.72** | **100** | **99.95** | **99.81** |
| **Grackle-2** | **99.95** | **99.95** | **99.95** | **99.95** | **99.95** | **99.95** | **99.95** | **99.95** | **99.95** | **99.95** | **99.77** | **99.77** | **99.91** | **99.81** | **99.67** | **99.95** | **100** | **99.77** |
| **Falcon** | **99.81** | **99.81** | **99.81** | **99.81** | **99.81** | **99.81** | **99.81** | **99.81** | **99.81** | **99.81** | **99.81** | **99.81** | **99.77** | **99.67** | **99.72** | **99.81** | **99.77** | **100** |

**Table S9.** Percentage identity of 18 PB1 segments

| **ID. for PB1** | **Bovine-1** | **Bovine-2** | **Cattle-1** | **Cattle-3** | **Cattle-4** | **Cattle-5** | **Cattle-6** | **Cattle-7** | **Feline-1** | **Feline-2** | **Human** | **Skunk** | **Blackbird-1** | **Blackbird-2** | **Goose** | **Grackle-1** | **Grackle-2** | **Falcon** |
| --- | --- | --- | --- | --- | --- | --- | --- | --- | --- | --- | --- | --- | --- | --- | --- | --- | --- | --- |
| **Bovine-1** | **100** | **99.96** | **100** | **100** | **100** | **100** | **99.96** | **100** | **100** | **100** | **99.82** | **99.82** | **99.96** | **99.96** | **99.82** | **100** | **100** | **99.96** |
| **Bovine-2** | **99.96** | **100** | **99.96** | **99.96** | **99.96** | **99.96** | **100** | **99.96** | **99.96** | **99.96** | **99.78** | **99.78** | **99.91** | **99.91** | **99.78** | **99.96** | **99.96** | **99.91** |
| **Cattle-1** | **100** | **99.96** | **100** | **100** | **100** | **100** | **99.96** | **100** | **100** | **100** | **99.82** | **99.82** | **99.96** | **99.96** | **99.82** | **100** | **100** | **99.96** |
| **Cattle-3** | **100** | **99.96** | **100** | **100** | **100** | **100** | **99.96** | **100** | **100** | **100** | **99.82** | **99.82** | **99.96** | **99.96** | **99.82** | **100** | **100** | **99.96** |
| **Cattle-4** | **100** | **99.96** | **100** | **100** | **100** | **100** | **99.96** | **100** | **100** | **100** | **99.82** | **99.82** | **99.96** | **99.96** | **99.82** | **100** | **100** | **99.96** |
| **Cattle-5** | **100** | **99.96** | **100** | **100** | **100** | **100** | **99.96** | **100** | **100** | **100** | **99.82** | **99.82** | **99.96** | **99.96** | **99.82** | **100** | **100** | **99.96** |
| **Cattle-6** | **99.96** | **100** | **99.96** | **99.96** | **99.96** | **99.96** | **100** | **99.96** | **99.96** | **99.96** | **99.78** | **99.78** | **99.91** | **99.91** | **99.78** | **99.96** | **99.96** | **99.91** |
| **Cattle-7** | **100** | **99.96** | **100** | **100** | **100** | **100** | **99.96** | **100** | **100** | **100** | **99.82** | **99.82** | **99.96** | **99.96** | **99.82** | **100** | **100** | **99.96** |
| **Feline-1** | **100** | **99.96** | **100** | **100** | **100** | **100** | **99.96** | **100** | **100** | **100** | **99.82** | **99.82** | **99.96** | **99.96** | **99.82** | **100** | **100** | **99.96** |
| **Feline-2** | **100** | **99.96** | **100** | **100** | **100** | **100** | **99.96** | **100** | **100** | **100** | **99.82** | **99.82** | **99.96** | **99.96** | **99.82** | **100** | **100** | **99.96** |
| **Human** | **99.82** | **99.78** | **99.82** | **99.82** | **99.82** | **99.82** | **99.78** | **99.82** | **99.82** | **99.82** | **100** | **99.65** | **99.78** | **99.78** | **99.65** | **99.82** | **99.82** | **99.78** |
| **Skunk** | **99.82** | **99.78** | **99.82** | **99.82** | **99.82** | **99.82** | **99.78** | **99.82** | **99.82** | **99.82** | **99.65** | **100** | **99.78** | **99.78** | **99.65** | **99.82** | **99.82** | **99.78** |
| **Blackbird-1** | **99.96** | **99.91** | **99.96** | **99.96** | **99.96** | **99.96** | **99.91** | **99.96** | **99.96** | **99.96** | **99.78** | **99.78** | **100** | **99.91** | **99.78** | **99.96** | **99.96** | **99.91** |
| **Blackbird-2** | **99.96** | **99.91** | **99.96** | **99.96** | **99.96** | **99.96** | **99.91** | **99.96** | **99.96** | **99.96** | **99.78** | **99.78** | **99.91** | **100** | **99.78** | **99.96** | **99.96** | **99.91** |
| **Goose** | **99.82** | **99.78** | **99.82** | **99.82** | **99.82** | **99.82** | **99.78** | **99.82** | **99.82** | **99.82** | **99.65** | **99.65** | **99.78** | **99.78** | **100** | **99.82** | **99.82** | **99.78** |
| **Grackle-1** | **100** | **99.96** | **100** | **100** | **100** | **100** | **99.96** | **100** | **100** | **100** | **99.82** | **99.82** | **99.96** | **99.96** | **99.82** | **100** | **100** | **99.96** |
| **Grackle-2** | **100** | **99.96** | **100** | **100** | **100** | **100** | **99.96** | **100** | **100** | **100** | **99.82** | **99.82** | **99.96** | **99.96** | **99.82** | **100** | **100** | **99.96** |
| **Falcon** | **99.96** | **99.91** | **99.96** | **99.96** | **99.96** | **99.96** | **99.91** | **99.96** | **99.96** | **99.96** | **99.78** | **99.78** | **99.91** | **99.91** | **99.78** | **99.96** | **99.96** | **100** |

**Table S10.** Percentage identity of 18 PB2 segments

| **ID. for PB2** | **Bovine-1** | **Bovine-2** | **Cattle-1** | **Cattle-3** | **Cattle-4** | **Cattle-5** | **Cattle-6** | **Cattle-7** | **Feline-1** | **Feline-2** | **Human** | **Skunk** | **Blackbird-1** | **Blackbird-2** | **Goose** | **Grackle-1** | **Grackle-2** | **Falcon** |
| --- | --- | --- | --- | --- | --- | --- | --- | --- | --- | --- | --- | --- | --- | --- | --- | --- | --- | --- |
| **Bovine-1** | **100** | **99.96** | **99.96** | **100** | **99.96** | **99.96** | **99.96** | **99.96** | **99.91** | **99.91** | **99.74** | **99.56** | **99.96** | **99.96** | **99.61** | **99.91** | **99.87** | **99.47** |
| **Bovine-2** | **99.96** | **100** | **100** | **99.96** | **100** | **100** | **100** | **100** | **99.96** | **99.96** | **99.78** | **99.61** | **100** | **100** | **99.65** | **99.96** | **99.91** | **99.52** |
| **Cattle-1** | **99.96** | **100** | **100** | **99.96** | **100** | **100** | **100** | **100** | **99.96** | **99.96** | **99.78** | **99.61** | **100** | **100** | **99.65** | **99.96** | **99.91** | **99.52** |
| **Cattle-3** | **100** | **99.96** | **99.96** | **100** | **99.96** | **99.96** | **99.96** | **99.96** | **99.91** | **99.91** | **99.74** | **99.56** | **99.96** | **99.96** | **99.61** | **99.91** | **99.87** | **99.47** |
| **Cattle-4** | **99.96** | **100** | **100** | **99.96** | **100** | **100** | **100** | **100** | **99.96** | **99.96** | **99.78** | **99.61** | **100** | **100** | **99.65** | **99.96** | **99.91** | **99.52** |
| **Cattle-5** | **99.96** | **100** | **100** | **99.96** | **100** | **100** | **100** | **100** | **99.96** | **99.96** | **99.78** | **99.61** | **100** | **100** | **99.65** | **99.96** | **99.91** | **99.52** |
| **Cattle-6** | **99.96** | **100** | **100** | **99.96** | **100** | **100** | **100** | **100** | **99.96** | **99.96** | **99.78** | **99.61** | **100** | **100** | **99.65** | **99.96** | **99.91** | **99.52** |
| **Cattle-7** | **99.96** | **100** | **100** | **99.96** | **100** | **100** | **100** | **100** | **99.96** | **99.96** | **99.78** | **99.61** | **100** | **100** | **99.65** | **99.96** | **99.91** | **99.52** |
| **Feline-1** | **99.91** | **99.96** | **99.96** | **99.91** | **99.96** | **99.96** | **99.96** | **99.96** | **100** | **100** | **99.74** | **99.56** | **99.96** | **99.96** | **99.61** | **99.91** | **99.87** | **99.47** |
| **Feline-2** | **99.91** | **99.96** | **99.96** | **99.91** | **99.96** | **99.96** | **99.96** | **99.96** | **100** | **100** | **99.74** | **99.56** | **99.96** | **99.96** | **99.61** | **99.91** | **99.87** | **99.47** |
| **Human** | **99.74** | **99.78** | **99.78** | **99.74** | **99.78** | **99.78** | **99.78** | **99.78** | **99.74** | **99.74** | **100** | **99.56** | **99.78** | **99.78** | **99.61** | **99.74** | **99.69** | **99.47** |
| **Skunk** | **99.56** | **99.61** | **99.61** | **99.56** | **99.61** | **99.61** | **99.61** | **99.61** | **99.56** | **99.56** | **99.56** | **100** | **99.61** | **99.61** | **99.52** | **99.56** | **99.52** | **99.39** |
| **Blackbird-1** | **99.96** | **100** | **100** | **99.96** | **100** | **100** | **100** | **100** | **99.96** | **99.96** | **99.78** | **99.61** | **100** | **100** | **99.65** | **99.96** | **99.91** | **99.52** |
| **Blackbird-2** | **99.96** | **100** | **100** | **99.96** | **100** | **100** | **100** | **100** | **99.96** | **99.96** | **99.78** | **99.61** | **100** | **100** | **99.65** | **99.96** | **99.91** | **99.52** |
| **Goose** | **99.61** | **99.65** | **99.65** | **99.61** | **99.65** | **99.65** | **99.65** | **99.65** | **99.61** | **99.61** | **99.61** | **99.52** | **99.65** | **99.65** | **100** | **99.61** | **99.56** | **99.69** |
| **Grackle-1** | **99.91** | **99.96** | **99.96** | **99.91** | **99.96** | **99.96** | **99.96** | **99.96** | **99.91** | **99.91** | **99.74** | **99.56** | **99.96** | **99.96** | **99.61** | **100** | **99.96** | **99.47** |
| **Grackle-2** | **99.87** | **99.91** | **99.91** | **99.87** | **99.91** | **99.91** | **99.91** | **99.91** | **99.87** | **99.87** | **99.69** | **99.52** | **99.91** | **99.91** | **99.56** | **99.96** | **100** | **99.43** |
| **Falcon** | **99.47** | **99.52** | **99.52** | **99.47** | **99.52** | **99.52** | **99.52** | **99.52** | **99.47** | **99.47** | **99.47** | **99.39** | **99.52** | **99.52** | **99.69** | **99.47** | **99.43** | **100** |
